# Single-cell whole-genome sequencing reveals convergent evolution in Burkitt lymphoma

**DOI:** 10.64898/2025.12.12.692552

**Authors:** Alexander S. Steemers, Markus J. van Roosmalen, Rico Hagelaar, Laurianne Trabut, Mark Verheul, Siem Jongsma, Jurrian K. de Kanter, Friederike Meyer-Wentrup, Ruben van Boxtel

## Abstract

Burkitt lymphoma (BL) carries multiple oncogenic drivers yet arises predominantly in children, whose normal cells harbour few age-related mutations. To investigate this paradox, we sought to define the sequence and timing of mutational drivers underlying BL development. Here, we analysed single-cell whole-genome sequencing (WGS) data of 250 paired normal and malignant B-cells from BL patients and integrated this data with 21 bulk WGS samples and an existing dataset of 157 clonally expanded B-cells from healthy individuals. Phylogenetic reconstruction revealed an accelerated accumulation of mutations following the emergence of the most recent common ancestor, leading to the early establishment of extensive genetic intra-tumoral heterogeneity. We further provide evidence that convergent evolution shapes this diversity at both the point mutation– and copy number variation–levels. This in-depth characterisation further establishes BL as a paradigmatic model of tumorigenesis with implications for therapeutic intervention.

## Introduction

Burkitt lymphoma (BL) has been termed the “Rosetta Stone” of cancer, reflecting its vital role in advancing our understanding of oncogenesis. It was the first cancer for which a viral association has been described (with the Epstein–Barr virus (EBV))^1^, the first tumour with a *MYC* chromosomal translocation^2^, and the first malignancy successfully treated with chemotherapy^3^. Despite these landmark discoveries, the mechanisms underlying BL initiation and clonal evolution, both within and across anatomical sites, remain poorly understood.

The genetic hallmark of BL is a reciprocal translocation that places the *MYC* oncogene under the control of immunoglobulin enhancers, resulting in constitutive MYC overexpression^4^. However, *MYC* dysregulation alone is insufficient for BL oncogenesis, and additional genetic alterations are required to fully drive malignant transformation^5–7^. These cooperating mutations typically involve genes important in B-cell receptor (BCR) signalling (*e.g.*, *ID3*, *FOXO1*, *TCF3*), cell survival and proliferation (*e.g.*, *MYC*, *CCND3*, *TP53*), and in SWI–SNF chromatin remodelling (*e.g.*, *SMARCA4*, *ARID1A*), highlighting recurrent disruption of signalling, growth regulation, and epigenetic networks in BL^8^.

Paradoxically, BL arises predominantly in children, posing the intriguing question of how multiple driver mutations accumulate within such a short developmental timeframe. This apparent contradiction highlights a critical gap in our understanding of BL evolution. Consequently, despite advances in defining the molecular drivers^7–11^, the timing and clonal architecture of BL mutations remain largely unresolved, in part because conventional bulk sequencing of biopsies blurs signals from genetically heterogeneous subclones.

Obtaining longitudinal tumour samples prior to diagnosis would provide invaluable insight into the clonal dynamics and the evolutionary steps preceding overt disease. In practice, however, such sampling is not possible. Consequently, most studies infer evolutionary history from single time-point samples, leveraging intra- tumoral heterogeneity (ITH) as a molecular record of past mutational events. In this study, we performed single-cell whole-genome sequencing (WGS) on both normal and malignant B-cells from primary BL samples, enabling direct tracing of clonal lineages and detailed characterization of ITH. By sampling tumour cells from either a single anatomical site or multiple spatially distinct regions within the same patient, we infer both the temporal and spatial evolution of BL.

## Results

### Single-cell WGS cohort represents broader BL mutational landscape

To reconstruct the clonal architecture and the sequence of genetic driver events, we performed single-cell WGS on six Burkitt lymphoma patients. These patients ranged in age from 4 to 17 years (median 13 years) and included five males and one female. Notably, two of these patients had multi-site involvement, allowing us to study dissemination dynamics. Specifically, two tumour site samples were collected at diagnosis for one patient, and paired diagnosis–relapse samples were obtained from another patient. Performing single-cell WGS on patient-matched normal and malignant B-cells requires the isolation of sufficient genomic DNA from individual cells, which is typically achieved via single-cell clonal expansion^12,13^. However, primary malignant B-cells are challenging to expand *ex vivo*, often requiring advanced culturing systems to support their survival and proliferation^14^. To circumvent this, we employed primary template-directed amplification (PTA) on flow cytometry-sorted single cells, enabling direct biochemical whole-genome amplification without the need for prior expansion^15^ (Fig. 1a and Supplementary Fig. 1).

**Fig 1.**
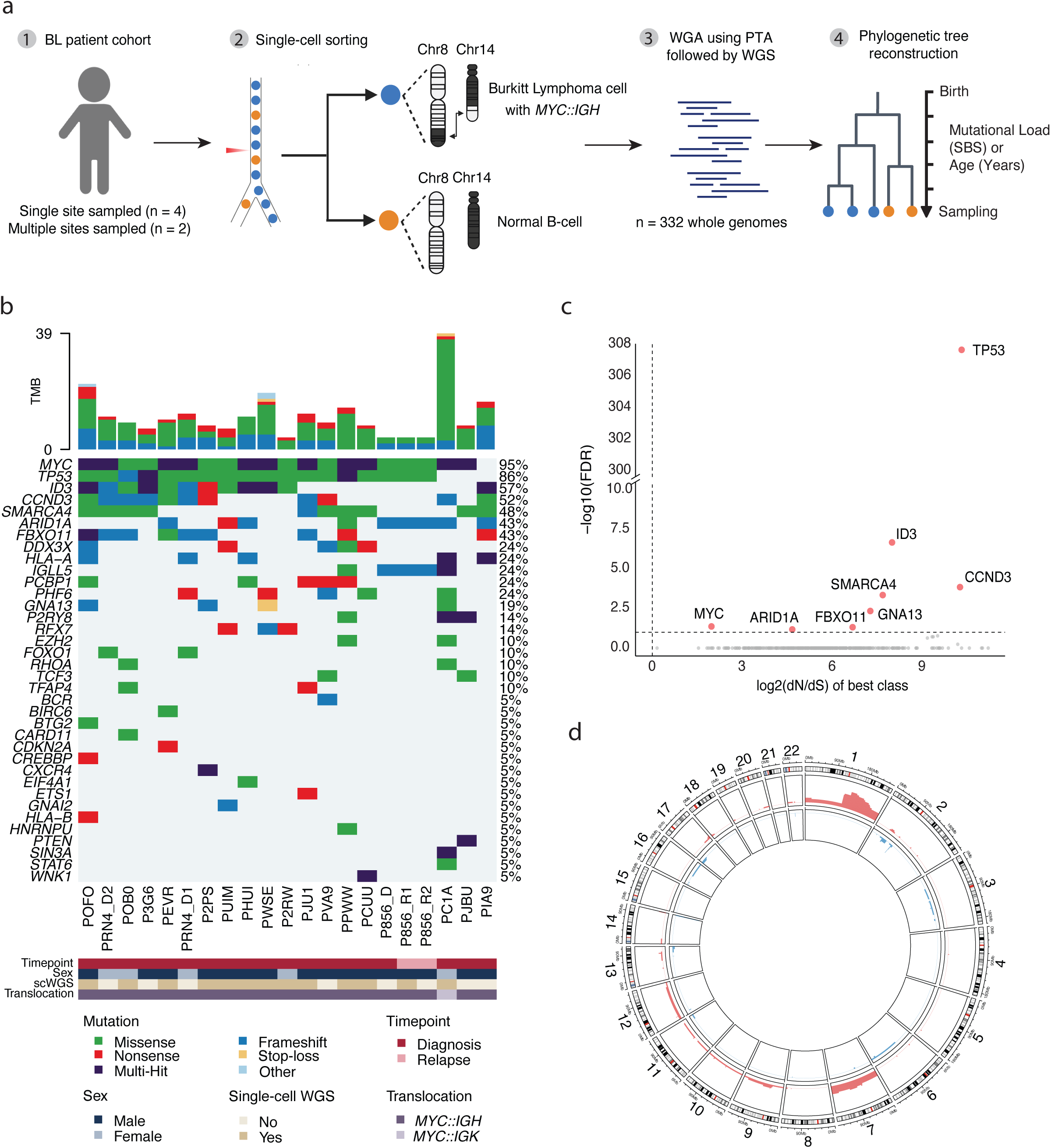
Burkitt lymphoma cohort and study design. **a**, Overview of the experimental design and schematic representation of the *MYC::IGH* (t(8;14)) translocation characteristic of Burkitt lymphoma cells. **b**, Oncoplot summarizing the landscape of SBS and INDEL driver mutations across 21 bulk WGS samples from 18 BL patients. Metadata below indicate sex, sampling timepoint, *MYC* translocation partner and inclusion in the single-cell WGS cohort. **c**, Gene-level positive selection analysis using dN/dS ratios. The x-axis shows the log₂(dN/dS) value of the best-fitting model class and the y-axis shows –log₁₀(FDR). Each point represents a gene, with significantly selected genes labelled. Dashed lines mark the significance (FDR < 10%) and neutrality (log₂(dN/dS) > 0) thresholds. **d**, Frequency of autosomal copy number gains (red) and losses (blue) across 18 BL patients.

To ensure that our single-cell WGS cohort is truly representative of BL, we additionally analysed bulk WGS data from a total of 18 BL patients (14 males, 4 females; median age 12 years, range 3–17). These comprised the 6 patients included in the single-cell WGS analysis and 12 additional cases (Supplementary Table 1). Assessment of viral DNA using *VIRUSBreakend*^16^, including EBV, led to no evidence of viral integration in any of the Burkitt samples (see Methods). This finding aligns with previous studies demonstrating that the majority of sporadic BL cases in children are EBV negative ^9,17^. Furthermore, this increases the biological uniformity of our cohort since it reduces any variability that may be introduced by an EBV infection. We next assessed the driver landscape of somatic mutations including single base substitutions (SBS), insertions and deletions (INDELs), and structural variants (SVs) (Fig. 1b and Supplementary Table 1).

The number of driver genes mutated per sample ranged from 4-13 (median = 7), reflecting the multi-step nature of BL tumorigenesis^8,18^. *MYC*, *TP53*, and *ID3* were the most frequently mutated genes, consistent with previous findings^7^. Notably, over half of *MYC*-mutant tumours (55%) and nearly half of *ID3*-mutant tumours (42%) harboured two or more mutations in the same gene, pointing towards strong selection pressure on these genes. However, with bulk WGS data, determining whether these mutations occur in the same or in distinct subclones remains inferential and cannot be confirmed with certainty^19^. To further elucidate the selection pressure, we applied dN/dS analysis to all somatic SBS and INDELs identified in our cohort (Fig. 1c). This method detects genes with an excess of nonsynonymous over synonymous substitutions, indicative of positive selection^20^. Indeed, all genes under significant positive selection (dN/dS > 1; FDR < 10%) were also among the most frequently mutated in our cohort, including *TP53*, *ID3*, *CCND3*, *MYC*, *SMARCA4*, *ARID1A*, *FBXO11*, and *GNA13*. In terms of copy number variations (CNVs), the most frequent gain was in chr1q (14/21 samples), in line with previous studies (Fig. 1d and Supplementary Table 1)^21,22^. Other recurrent events included gains in chr7q (9/21), chr7p (4/21), and chr1p (3/21). Taken together, our single-cell WGS cohort closely represents the genetic landscape of BL.

### Phylogenetic reconstruction

Following single-cell WGS, 82 out of 332 single cells were excluded due to low sequencing coverage or allelic imbalance (see Methods), resulting in a final dataset of 250 high-quality single-cell genomes (Supplementary Table 2). A mean of 42 whole genomes were taken forward per patient (range 16-58 whole genomes), with a mean sequencing depth of 11.7X (range 7.6-24.6X) and an average of 83% genome coverage at ≥ 5X depth. Following germline and artefact filtering, we identified a total of 283,400 autosomal SBS and 51,164 autosomal INDELs^23^. To explore the distribution of these mutations between the different cells of the same patient, we reconstructed phylogenies based on shared genome-wide SBS, enabling us to resolve the clonal relationships at single-cell resolution (Fig. 2). In these trees, internal branches represent mutations propagated across multiple descendants from a shared ancestral cell, terminal branches capture more recent mutations unique to individual cells, and nodes mark inferred ancestral cells from which downstream lineages arose, offering a unique window into the past. All trees displayed a dense cluster of coalescences (branching nodes) near the root, reflecting the rapid expansion of B-cell precursors and the resulting genomic divergence during early lymphopoiesis, similarly as previously reported for healthy haematopoiesis^24,25^. This indicates that for any two normal B-cells, their most recent common ancestor (MRCA) lies early in development. Beyond this early expansion, the phylogenies of normal B-cells displayed a ‘comb-like’ branching pattern. However, in one patient (PIA9), the normal B-cell phylogeny deviated from this expected structure and instead showed evidence of clonal expansion within the normal B-cell compartment (Fig. 2). Subsequent driver analysis revealed that the clonally expanded normal B- cell population uniquely harboured a missense mutation in *ACTG1*, a gene previously reported to be under positive selection in normal B-cells from healthy donors^13^.

**Fig 2.**
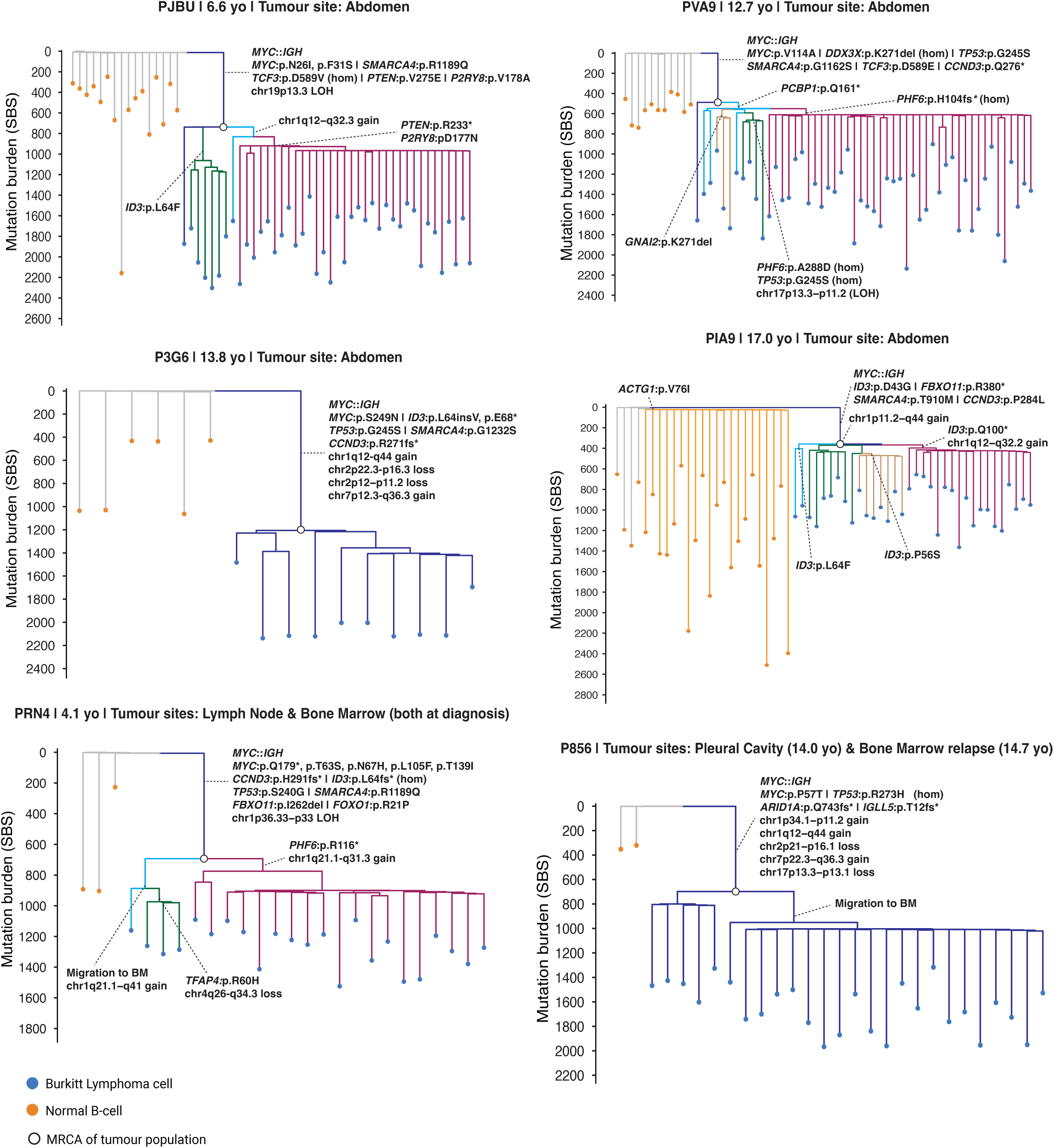
Phylogenetic trees of Burkitt lymphoma patients. Each phylogenetic tree shows the pattern of shared somatic single base substitutions (SBS) among sequenced single cells. Branch lengths are proportional to mutation counts shown on the vertical axes. Private branches represent those mutations unique to a single cell, whereas shared branches represent mutations found in two or more cells. Branching points (coalescences) correspond to ancestral cells that underwent symmetrical self-renewing divisions, where descendants of both daughter cells were sampled. These points mark the most recent common ancestor (MRCA) of the downstream clade sharing that lineage. Dots at the bottom represent individual sequenced cells, coloured according to cell type: blue for Burkitt lymphoma cells and orange for normal B-cells. The MRCA of all tumour cells for each patient is depicted with a black circle. Branches containing different driver mutations and copy number alterations are highlighted by colour. Only drivers that occurred in at least two cells are annotated. Dissemination of tumour cells to different anatomical sites is also annotated for patients PRN4 and P856.

### BL cells exhibit convergent evolution

A distinctive feature of the tumour phylogenies is the clustering of malignant cells into a single clade, marked by a long trunk. B-cell receptor (BCR) recombination analysis showed that all tumour cells had the same immunoglobulin heavy-light chain rearrangement, confirming that the clone started expanding following this process (i.e. monoclonal). In contrast, normal B-cells remained polyclonal (Supplementary Fig. 2). This indicates that in the case of BL cells, the identity of the MRCA was a B-cell, whereas in the case of normal B-cells it is a more primitive cell. In line with previous reports describing a high frequency of BCR rearrangements involving the IGHV3 genes^10,26^, this V gene family was observed in 5/6 of the tumours. When comparing each patient’s bulk tumour sample to the corresponding phylogenetic tree, 93.2% of truncal mutations, 46.9% of intermediate mutations, and 0.1% of private branch mutations were detected (Supplementary Fig. 3). This does not only demonstrate the high recall rate of our single-cell WGS approach but also underscores the limitations of bulk sequencing in capturing subclonal and private variants.

Driver analysis revealed that the hallmark *MYC::IGH* translocation was always located in the shared trunk of the tumour phylogenies and was consistently accompanied by additional truncal mutations. Unexpectedly, we were able to place known driver mutations in the intermediate branches of the trees. These intermediate nodes seeded rapidly expanding subclones, as evidenced by bursts of coalescences emerging directly beneath them. Although we could not place any known driver mutations in the intermediate branches of patient P3G6, subclonal expansions were still observed. This observation indicates that BL tumours exhibit substantial ITH, where multiple clonal lineages arise from a common ancestor. Within these subclonal driver mutations, we found strong evidence of convergent evolution. This occurs when separate tumour lineages independently acquire similar genetic alterations, including mutations in the same driver genes or CNVs affecting the same chromosomal regions. Specifically, in PJBU, both *PTEN* and *P2RY8* were independently mutated in the trunk and in an intermediate branch that led to the largest subclone; in PVA9, *PHF6* was recurrently altered across two separate intermediate branches; and in PIA9, three distinct *ID3* mutations arose in intermediate lineages, each driving parallel subclonal expansions. In addition to SBSs and INDELs, we found evidence of convergent evolution at the CNV level, particularly affecting chromosome 1. In PIA9, two independent gains were detected (chr1p11.2–q44 and chr1q12–q32.2), while PRN4 exhibited a similar pattern, with a chr1q21.1–q41 gain in the bone marrow and a chr1q21.1–q31.3 gain in the lymph node. These findings demonstrate that tumour evolution is not purely stochastic but instead shaped by convergent selective pressures that repeatedly target the same genomic regions and pathways, driving parallel subclonal trajectories.

Our single-cell WGS cohort also included two patients with dissemination, allowing us to dissect the spatial evolution of tumours. In PRN4, an ancestral clone appeared to bifurcate. One branch migrated to the bone marrow and subsequently acquired a site-specific subclonal *TFAP4* mutation, while the other branch presumably remained in the lymph node and acquired an additional *PHF6* mutation. This pattern demonstrates that ITH does not only exist across, but also within anatomical sites. In P856, no subclonal driver mutations were found in either the diagnostic or relapse samples, indicating that the tumour expansion in both diagnosis and relapse was not fuelled by the acquisition of additional genetic drivers. Instead, the phylogeny was characterized by a single dominant truncal lineage. A previous study has indicated that relapse patients can be characterized by having complex karyotypes and being *TP53* deficient due to two mutations, namely an SBS variant and a low-frequency CNV^27^. Indeed, patient P856 acquired multiple CNVs across the genome and a homozygous R273H mutation in *TP53* due to a missense mutation combined with a chr17p13.3-13.1 focal loss. All other patients in our cohort who did not relapse displayed a heterozygous clonal *TP53* mutation, with the exception of PVA9, where a subclonal homozygous G245S *TP53* mutation arose due to a local LOH. However, this alteration was present only in a minor subclone within the tumour mass, suggesting that the timing of a driver event may be more critical than the mutation itself in determining tumour aggressiveness.

### Copy-number variations represent late events in tumour evolution

Our single-cell approach enabled us to resolve the exact sequential order of CNV events. Strikingly, many CNVs presented as subclonal events, include chromosome 1 gains in PJBU, PIA9 and PRN4, a chromosome 17p LOH in PVA9, and a chromosome 4q loss in only a subset of tumour cells in the bone marrow of PRN4. To validate the late timing of CNVs in a larger cohort, we performed a relative timing analysis on our bulk WGS data (see Methods). This approach estimates timing in regions with copy number gains or loss of heterozygosity (LOH) by considering the co-occurrence of SBS and INDELs. Specifically, the relative timing of a CNV is calculated as the ratio of amplified to non-amplified mutations within the affected region. In line with our single-cell data, we found that CNV timings were not evenly distributed, instead, they tended to arise late (Supplementary Fig. 4a). Notably, the recurrent copy number gains mentioned earlier in our BL cohort (chr1p, chr1q, chr7p and chr7q) consistently showed a late relative timing (Supplementary Fig. 4b). Taken together, our results demonstrate that CNVs represent one of the late genomic events preceding the establishment of the MRCA, and in many cases are acquired as subclonal events.

### Accelerated mutation rate in BL cells

We next sought to compare the SBS mutational burden between normal and BL cells. It was previously reported that bulk tumour samples harbour the same mutational burden as age-matched normal memory B-cells^13^. To validate our observations within a larger dataset, we incorporated 157 previously clonally expanded single-cell genomes of naïve and memory B-cells from 7 healthy donors^13^. Since SBS9 is associated with somatic hypermutation and therefore distinguishes naïve B-cells from differentiated memory B-cells^13^, we subdivided our patient-derived normal B-cells into SBS9⁺ and SBS9⁻. We then applied a linear mixed-effects model trained on naïve B-cells from healthy donors to estimate the expected mutation load at each age, thereby providing an age-adjusted observed-vs-expected (O/E) ratio baseline (Fig. 3a). Patient-derived SBS9⁻ normal B-cells (median O/E ratio 1.49) displayed slightly higher mutation load compared to naïve B-cells (median O/E ratio 0.93) from healthy donors. Further analysis showed that this was skewed due to the SBS9⁻ normal B-cells from patient PIA9 which contributed to a higher mutation load (Supplementary Fig. 5). By contrast, memory B-cells from healthy donors and SBS9⁺ normal B-cells from patients both exhibited elevated mutation burden, with no significant difference between them (median O/E ratio 1.97 vs 2.36; p-value = 0.59). Likewise, no significant difference was observed when comparing SBS9⁺ normal B-cells to the clonal mutations identified in our bulk WGS tumour samples (median O/E ratio 2.48; p-value = 0.22), as previously reported^13^. This indicates that the mutation rate in the lineage that eventually gave rise to the BL was similar as normal B-cells up to the point of the MRCA. Importantly, single malignant cells showed a marked increase in mutation load compared to SBS9⁺ normal B-cells, with up to a 7.42-fold increase in mutational load compared to baseline, indicating that the mutation rate increased after the Burkitt lymphoma MRCA started expanding.

**Fig 3.**
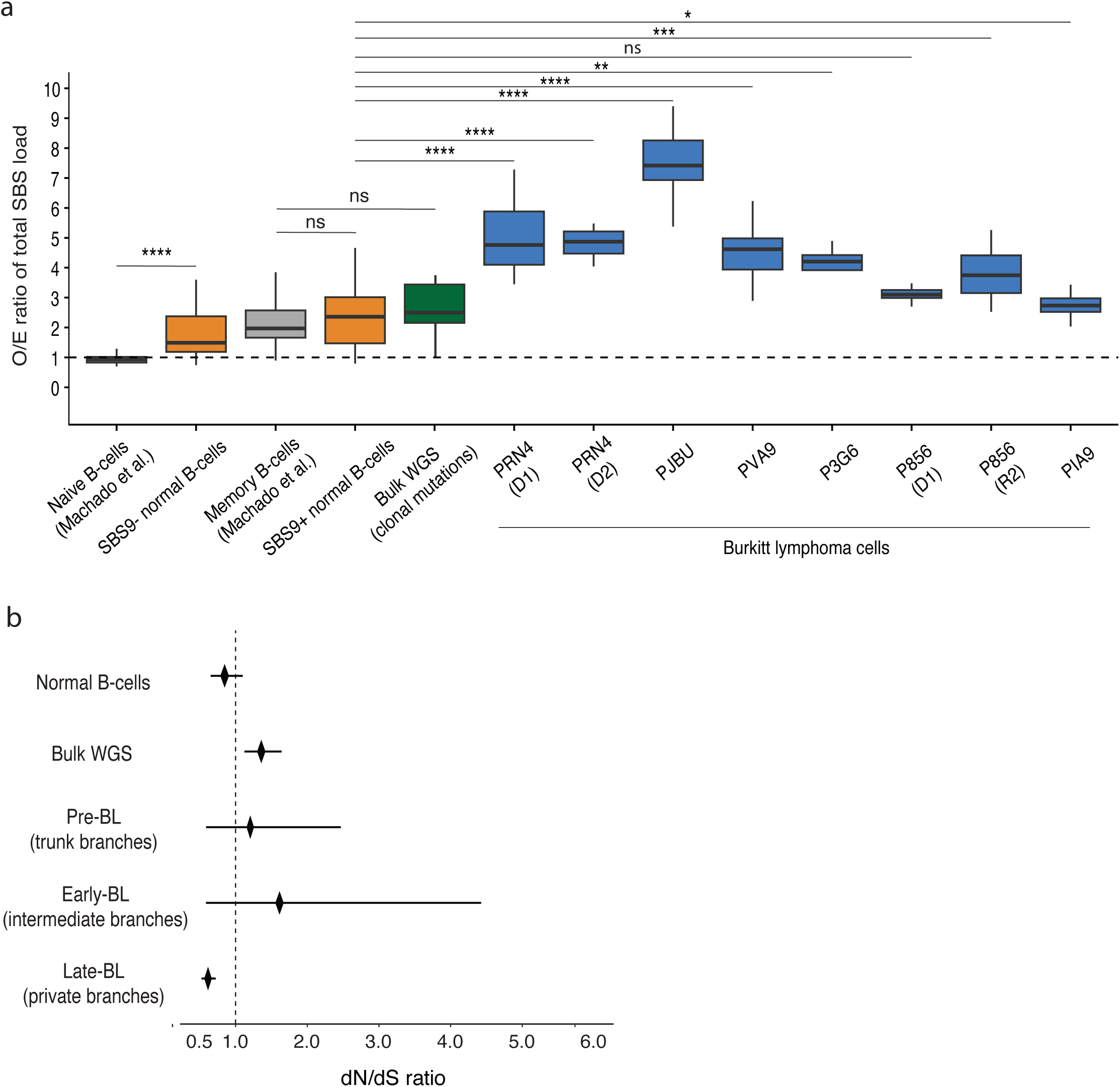
Somatic mutation burden and selection dynamics in BL cells. **a**, Observed/expected (O/E) ratio of somatic SBS load across single and bulk B-cell populations. Naïve and memory B-cells derived from healthy donors (Machado et al.) are shown in grey, SBS9⁺ and SBS9⁻ B-cells derived from BL patients in orange, clonal mutations from BL bulk WGS samples in green, and tumour cells from BL patients in blue. The dashed line indicates the expected SBS load (O/E = 1), estimated from a linear mixed-effects model fitted to naïve B-cells. Statistical significance was assessed using a Wilcoxon rank-sum test comparing BL cells from each patient to normal SBS9⁺ B-cells. Significance levels are denoted as P < 0.05 (*), P < 0.01 (**), P < 0.001 (***), and P < 0.0001 (****). **b**, dN/dS ratios across normal B-cells and distinct evolutionary stages of Burkitt lymphoma. Diamonds show the maximum-likelihood estimate (MLE) of the overall dN/dS ratio per group with 95% CIs (horizontal bars). Ratios were separately calculated for different stages of BL evolution: pre-BL (trunk branches) representing mutations acquired before the MRCA, early-BL (intermediate branches) representing mutations accumulated prior to major clonal expansions, and late-BL (private branches) representing mutations acquired after clonal expansions. The dashed vertical line marks neutrality (dN/dS = 1). Immunoglobulin loci (IGH, IGK, IGL) were excluded prior to fitting.

Having established that mutation rates accelerate after the MRCA, we next wondered what the consequence of this is on selection. For this, we performed dN/dS analysis across normal B-cells and distinct phases of Burkitt lymphoma evolution (Fig. 3c). Normal B-cells exhibited dN/dS values close to neutrality (dN/dS = 0.85, 95% CI 0.65–1.10). In contrast, both the pre-expansion phase of BL, i.e. trunk branches, (dN/dS = 1.40, 95% CI 0.82–2.39) and the early-BL phase, i.e. intermediate branches (dN/dS = 1.95, 95% CI 0.72–5.27) showed dN/dS estimates > 1, consistent with positive selection. To confirm this in a larger cohort, we also looked at the dN/dS ratio of the 21 bulk WGS samples which, as we have shown earlier, largely consist of pre-expansion and early-BL mutations (Supplementary Fig. 3). In line with our single-cell data, these mutations yielded a dN/dS ratio of 1.36 (95% CI 1.12-1.64), consistent with positive selection. Surprisingly, the private branches displayed evidence of purifying selection (dN/dS = 0.62, CI 0.52–0.72), indicating that once major drivers had been acquired, further nonsynonymous mutations might harm the fitness of the clones and are thus selectively constrained. Collectively, these findings support a model in which Burkitt lymphoma originates from pre-malignant cells with normal mutation rates under early positive selection, transitions at the MRCA into an accelerated mutational phase marked by continued positive selection and acquisition of additional driver mutations, and subsequently undergoes clonal expansions constrained by purifying selection.

### Spatial and temporal dynamics of mutational processes

We next wondered which mutational processes lead to the excess mutational load associated with malignant outgrowth. For this, we investigated the mutational processes active in both normal and malignant B-cells by quantifying mutational signature contributions across individual branches of the phylogenetic trees (Fig. 4a and Supplementary Fig. 6 and 7). Mutational signature analysis extracted the following single bases substitution signatures: SBS1, SBSblood, SBS7a, SBS17b, SBS9, and SBS18 (see Methods). SBS1, caused by spontaneous deamination of methylated CpGs and linked to replication^28,29^, and SBSblood, thought to be a hematopoietic form of the SBS5 process^13^, are both clock-like endogenous signatures. SBS7a and SBS17b, on the other hand, are both caused by exogenous processes, since SBS7a is characteristic of ultraviolet light exposure and SBS17b is believed to be caused by a specific microenvironmental exposure associated with tissue residency in gastrointestinal mucosa^13,30^. As mentioned earlier, SBS9 is attributed to somatic hypermutation while SBS18 is linked to damage caused by reactive oxygen species^31^. No additional mutational signatures were found in BL cells compared to normal B-cells, suggesting that existing mutational processes are responsible for the increased mutation load.

**Fig 4.**
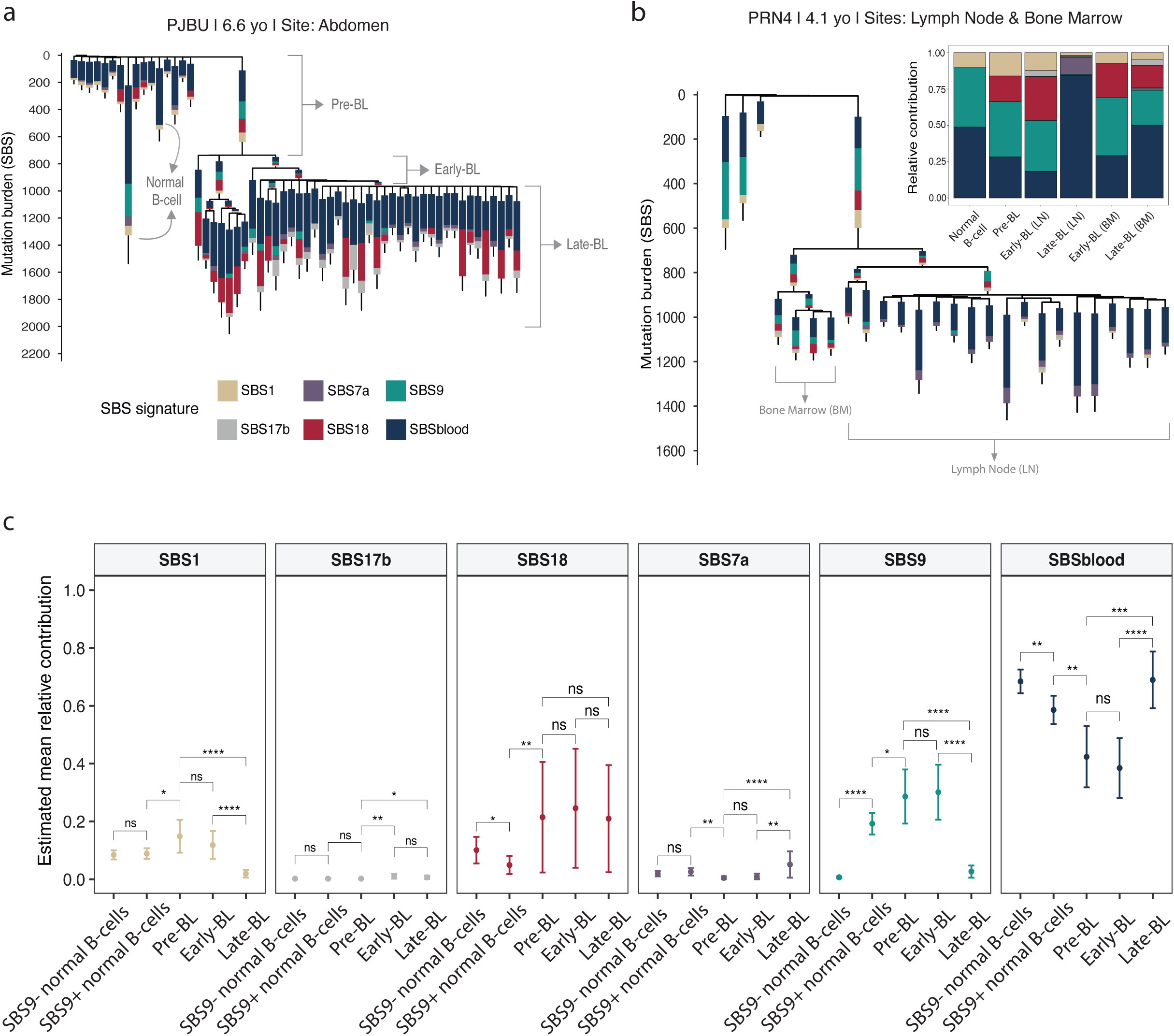
Mutational processes active during BL tumorigenesis. **a**, A phylogenetic tree for an example patient PJBU with the estimated proportion of SBS mutations attributed to SBS signatures SBS1, SBS7a, SBS9, SBS17b, SBS18 and SBSblood. Branches were subdivided into four categories: (1) branches of MYC::IGH-negative (normal B-cells), (2) ancestral branch of MYC::IGH-positive cells representing the lineage of BL origin (‘pre-BL’), (3) intermediate branches representing early mutations after the MRCA (‘early-BL’), and (4) private branches of MYC::IGH-positive BL cells representing mutations acquired after clonal expansions (‘late-BL’). **b**, The phylogenetic tree for patient PRN4 and the relative contribution of SBS signatures for each branch category, with the added information of the anatomical site from which tumour cells were sampled. **c**, The estimated mean relative contribution of SBS signatures in each branch category. Normal B-cells were further divided into SBS9⁻ and SBS9⁺. Statistical significance was assessed using a two-sided Wald z-test. Significance levels are denoted as P < 0.05 (*), P < 0.01 (**), P < 0.001 (***), and P < 0.0001 (****).

In patient PRN4, from which multiple sites were sampled at diagnosis, both lymph node and bone marrow tumours shared a common ancestral branch dominated by endogenous processes, namely SBS1, SBSblood, and SBS9 (Fig. 4b). However, site-specific private branches revealed distinct mutational exposures. Private mutations in lymph node–derived tumour subclones showed a high contribution from SBS7a (relative SBS7a contribution = 14.1%), which was not observed in the bone marrow–derived tumour subclones (relative SBS7a contribution = 1.6%). This is consistent with the anatomical location of the affected lymph node in the head and neck region, which could plausibly be exposed to ultraviolet light mutagenesis. These findings suggest that local microenvironmental or tissue-specific exposures may shape the mutational landscape of spatially separated Burkitt lymphoma cells.

We next wanted to assess the temporal dynamics of mutational processes. SBS9 remained active in early-BL branches but dropped sharply in late-BL branches (Fig. 4c). This might suggest that SBS9 plays an important role in driving the positive selection pressure during the early stages of malignant transformation but is no longer active when tumour cells start clonally expanding. Instead, the relative contribution of SBSblood increased in late-BL branches compared to pre-BL and early-BL branches, indicating that this endogenous process becomes more prominent during the later stages of malignant evolution and is associated with the increased mutation rate post-MRCA expansion.

### Timing BL expansion

We further sought to time the driving events and clonal expansions in years. To convert our phylogenetic trees from somatic SBS mutations to time, we first determined which processes accumulate at a constant mutation rate, as not all processes are expected to be constantly active with age. Hence, mutational signature contributions were examined in normal cells and correlated with donor age at the timepoint of sampling (Fig. 5a). SBS1 (slope = 1.1 SBS/year, p = 0.0464) and SBSblood (slope = 22.1 SBS/year, p = 5.71x10^-4^) both correlated linearly with time, as previously shown^13^. In contrast, SBS7a (p = 0.465) and SBS9 (p = 0.511) did not correlate significantly with age, and in many normal cells these signatures were absent. SBS17b showed no detectable contribution in normal cells. Although SBS18 exhibited a strong positive correlation with age (slope = 15 SBS/year, p = 7.4x10^-5^), many normal cells again lacked any detectable contribution from this signature. Altogether, these observations suggest that SBS1 and SBSblood indeed represent constant clock-like endogenous mutational processes, while SBS7a, SBS9, SBS17b and SBS18 likely arise through more stochastic or episodic processes^13^. As a result, we subset our phylogenetic trees to only include mutations attributed to SBS1 and SBSblood, and used those trees to calibrate branch lengths in years (see Methods).

**Fig. 5.**
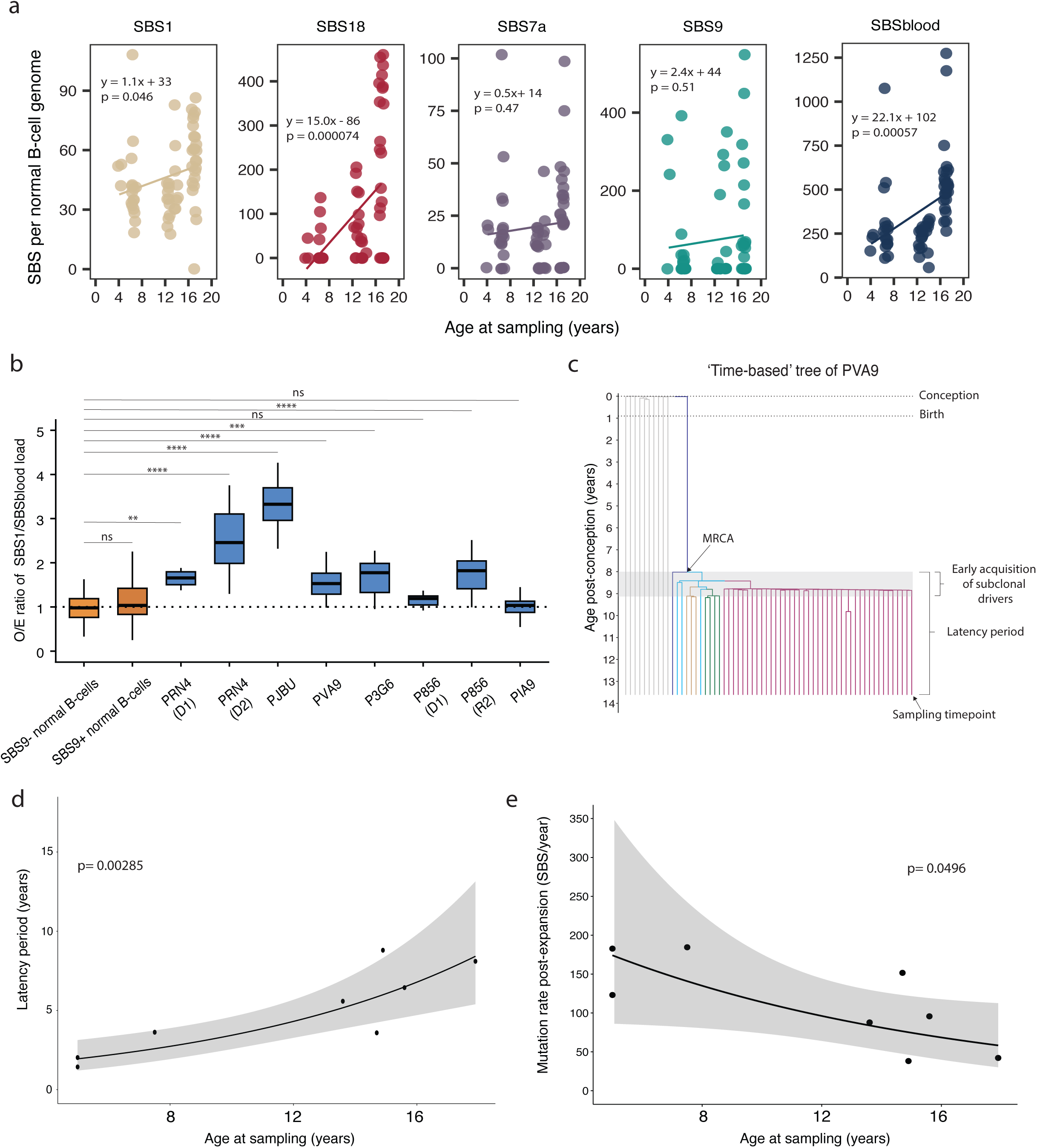
Timing of BL clonal expansion and dissemination. **a**, SBS mutation burden per normal B-cell genome, separated by mutational signature. Lines represent the fitted values from linear mixed-effects models for each signature. Signature SBS17b showed no detectable contribution in any normal B-cell and is therefore not displayed. **b**, Observed/expected (O/E) ratio of somatic SBS1/SBSblood mutation load across normal and malignant B-cells. SBS9+ and SBS9- B-cells are shown in orange, while tumour cells are shown in blue. The dashed line indicates the expected load (O/E = 1), estimated from a linear mixed-effects model fitted to SBS9⁻ normal B-cells. Statistical significance was assessed using a Wilcoxon rank-sum test comparing normal SBS9+ and BL cells from each patient to normal SBS9⁻ B-cells. Significance levels are denoted as P < 0.05 (*), P < 0.01 (**), P < 0.001 (***), and P < 0.0001 (****). **c**, ‘Time-based’ tree of patient PVA9, in years post-conception. Each clade that acquired an additional subclonal driver mutation is highlighted in a distinct colour to illustrate independent evolutionary trajectories within the tumour. **d**, Latency period (time from MRCA emergence to diagnosis) plotted against age at sampling. Each point represents one tumour sample. An exponential model was fitted (solid line) with 95% confidence interval (shaded area). **e**, Mutation rate post-expansion (i.e. mean mutation load per tumour cell post-MRCA divided by the latency period) plotted against age at sampling. Each point represents one tumour sample. An exponential model was fitted (solid line) with 95% confidence interval (shaded area).

After restricting our analysis to SBS1 and SBSblood, we found that both normal B-cell subtypes (SBS9⁻ and SBS9⁺) exhibited O/E ratios close to one, implying that these processes indeed accumulate mutations at a comparable rate despite cell type (Fig. 5b). Nevertheless, malignant cells displayed a higher O/E ratio, with a notable exception in the diagnostic samples of P856 and PIA9. This indicates an acceleration of these otherwise stable, clock-like mutational processes after malignant transformation, which could be due to either an increased cellular turnover or a decreased DNA repair efficiency. To test the latter, we analysed transcriptional strand bias and observed no change in the late private mutations, suggesting that the transcription-coupled repair pathway is not impaired during late tumour evolution (Supplementary Fig. 8). Based on these findings we inferred the timing of the coalescences using the *rtreefit* package^32^ while accounting for a higher rate of mutation accumulation at the end of the trunk branch of the tumour clade, just before the emergence of the MRCA.

From our ‘time-based’ trees, we could now time several key events, including the emergence of the MRCA, the acquisition of subclonal drivers, and the latency period, which is the time between the MRCA of the tumour and sampling (Fig. 5c). Despite the clinically aggressive nature of Burkitt lymphoma, our data indicate latency periods of several years (mean 4.95 years; range 1.44 – 8.80 years). Interestingly, we were able to time the origin of the relapse in patient P856 ∼6 years before the initial diagnosis (Supplementary Fig. 9). Although our cohort was small, there seemed to be a trend where younger patients correlated with a shorter latency period and faster post-expansion mutational rates (Fig. 5d, e). This has previously also been described in *BCR::ABL1*-driven chronic myeloid leukaemia, where younger patients showed more explosive growth together with a shorter duration between the beginning of *BCR::ABL1* clonal expansion and diagnosis^33^. Moreover, subclonal drivers were acquired soon after the establishment of the MRCA (1-1.5 years post-MRCA), suggesting that ITH arises during early tumour development followed by stable expansion of a limited number of dominant clones (Supplementary Fig. 9).

## Discussion

In this study, we leveraged the high resolution of single-cell WGS to resolve the ITH and reconstruct the temporal and spatial sequence of genetic events in BL. We found that BL cells harbour a substantially higher mutational burden than age-matched normal B-cells. This contrasts with previous bulk sequencing studies that reported comparable mutation loads between BL and normal B-lymphocytes^13^, underscoring the greater sensitivity of single-cell WGS for capturing genetic diversity.

Reconstruction of phylogenetic trees revealed that clonal expansions were driven by subclonal, fitness-enhancing mutations arising after the MRCA. This suggests that the hallmark *MYC::IGH* translocation and other early genetic drivers are important in enabling tumour initiation and survival, but do not directly account for the explosive growth of BL. We found compelling evidence of convergent evolution, where distinct subclones independently acquired mutations in the same genes or CNVs affecting the same chromosome. dN/dS analysis across phylogenetic branches revealed strong positive selection during the pre-malignant and early tumour evolution, consistent with acquisition of key driver mutations. Following major clonal expansions, selection shifted toward a purifying regime, likely reflecting spatial constraints and competition for resources within the tumour microenvironment. In line with this, SBS9 activity, associated with DNA polymerase η–mediated mutagenesis, was confined to early evolutionary branches, suggesting it may contribute to early driver acquisition and positive selection.

Our cohort also included a patient sampled at diagnosis and relapse. In this case, phylogenetic reconstruction revealed a linear evolutionary trajectory in which all driver mutations were present in the ancestral clone, with no evidence of subclonal diversification. This implies that tumour expansions in each timepoint were likely driven by non-genetic or microenvironmental factors. Notably, this patient harboured a clonal homozygous *TP53* mutation caused by a missense mutation and local copy-number loss; a similar event in another patient (PVA9) was subclonal. This suggests that the timing rather than the presence of *TP53* mutations may determine their clonal dominance^34^.

Temporal calibration of phylogenetic trees showed that the MRCA emerged several years before diagnosis (mean 4.95 years), with latency period positively correlating with patient age. This finding raises important questions about which cell-intrinsic and extrinsic factors sustain prolonged latency and, conversely, what triggers the rapid clinical progression characteristic of BL. Moreover, our temporal analysis showed that subclonal driver mutations were acquired within 1–1.5 years of MRCA emergence, indicating that ITH arises early rather than accumulating gradually during tumour growth.

Collectively, our results demonstrate that BL tumours are characterized by pronounced intratumoral heterogeneity which is shaped by convergent evolution both on the small-scale (SBSs and INDELs) and large-scale (CNVs) genomic levels. These observations imply that therapeutic strategies targeting subclonal pathways sustaining tumour proliferation may be more effective than interventions solely directed at early events such as *MYC* activation, which have yet to show clinical efficacy.

## Methods

### Patient samples

Samples were collected from patients with Burkitt lymphoma via the biobank of the Princess Máxima Center for Pediatric Oncology in accordance with the Declaration of Helsinki. Patients provided informed, written consent for the use of their samples for research. For each sample, viably frozen single-cell suspensions were obtained, which is part of our routine diagnostic workup ^35^. In addition, bulk WGS data (tumour and matched normal samples) from BL patients was obtained from our in-house diagnostics database, where samples are routinely sequenced as part of clinical evaluation. The study was covered under the proposal PMCLAB2022-303 approved by the Institutional Review Board of the Máxima. Clinical information of the patients, including which sites where sampled, is summarized in Supplementary Table 1.

### Sample work-up

Samples were worked up for single-cell WGS according to our previously published STAR Protocol guidelines^36^. Briefly, samples were stained for fluorescence-activated cell sorting (FACS) after thawing. Burkitt lymphoma (BL) cells and normal B-cells were purified on a SH800S Cell Sorter (Sony) using a 100 μm microfluidic sorting chip. First, manual gates were used to remove debris (FSC-A vs. SSC-A), doublets (FSC-A vs. FSC-H) and dead cells (DAPI+). BL cells were identified using the following surface markers: CD20+CD10+ and either IgK+ or IgL+ (depending on their monoclonal immunoglobulin light chain rearrangement). Single BL cells and normal B-cells (CD20+) were index sorted in a 96-well plate or 384-well plate prepared with PTA-buffer. Bulk BL cells were then sorted for DNA isolation. Representative gating strategies are shown in Supplementary Fig. 1. All antibodies were obtained from BioLegend. Antibodies used for BL and normal B-cell populations: CD20-FITC (clone 2H7, 1:25, #302303), CD10-APC (clone HI10a, 1:50, #312209), IgK-PE/Cy7 (clone MHK-49, 1:50, #316520), IgL-AF700 (clone MHL-38, 1:25, #316631).

### Germline controls

Mesenchymal stromal cells (MSCs) were cultured from the bone marrow fraction of 500,00 cells/well in 12-well culture dishes with 2 mL Advanced DMEM-F12 medium (#12634010, Gibco) supplemented with 10% FBS (Gibco), 1% GlutaMax (#35050061, Gibco) and 1% Penicillin-Streptomycin (#15140122, Thermo Fisher Scientific). Medium was refreshed every other day to remove non-adherent cells and MSCs were be harvested when confluent (after approximately 3 weeks).

### DNA isolation and WGS

DNA was isolated from cell pellets of bulk BL cells and MSCs using the DNeasy DNA Micro Kit (#56304, Qiagen), following the manufacturer’s instructions. The standard protocol was slightly adjusted by adding 2 µL RNase A (#19101, Qiagen) during the lysis step and eluting DNA in 50 µL low EDTA TE buffer (10 mM Tris, 0.1 mM EDTA, G Biosciences, #786150).

DNA from single cells was amplified using the ResolveDNA® WholeGenome Amplification Kit (#100136, BioSkryb) or the ResolveDNA® Whole Genome Amplification Kit v.2.0 (#100545, BioSkryb) using a D100 Single Cell Dispenser (HP), according to the manufacturer’s instructions. Details on which kit was used per sample is summarized in Supplementary Table 2.

For each sample, DNA libraries for Illumina sequencing were generated from at least 45 ng genomic DNA using standard protocols. For PTA-amplified DNA at least 200 ng genomic DNA was used. The libraries were sequenced at a depth of 15x (single cells) or 30x (bulk tumour and MSC samples).

### Read mapping

Sequencing reads were first mapped to genome GRCh38 using the BWA-MEM2 (Burrows–Wheeler Aligner) algorithm (v.2.2.1) using the settings “–M –c 100 -R”. Duplicates were then marked using GATK4spark (v.4.4.0.0), and base recalibration was performed using GATK4 (v.4.4.0.0).

### Mutation calling, filtering and annotation

Mutation calling was performed using GATK’s HaplotypeCaller on combined samples per patient. Variant quality score recalibration (VQSR) was applied separately for SNPs and INDELs using the VariantRecalibrator tool with the following annotations: -an QD -an MQ -an MQRankSum -an ReadPosRankSum -an FS -an SOR -mode SNP for SNPs, and -an QD -an MQ -an MQRankSum -an ReadPosRankSum -an FS -an SOR -mode INDEL for INDELs. ApplyVQSR was then run with --truth-sensitivity-filter-level 99.9 -mode SNP and --truth-sensitivity-filter-level 99.9 -mode INDEL.

Next, GATK’s VariantFiltration was run with the following options: “--filter-expression ’QD < 2.0’ --filter-expression ’MQ < 40.0’ --filter-expression ’FS > 60.0’ --filter-expression ’HaplotypeScore > 13.0’ --filter-expression ’MQRankSum < -12.5’ --filter-expression ’ReadPosRankSum < -8.0’ --filter-expression ’MQ0 >= 4 CC ((MQ0 / (1.0 * DP)) > 0.1)’ --filter-expression ’DP < 5’ --filter-expression ’QUAL < 30’ --filter-expression ’QUAL >= 30.0 CC QUAL < 50.0’ --filter-expression ’SOR > 4.0’ --filter-name ’SNP_LowQualityDepth’ --filter-name ’SNP_MappingQuality’ -- filter-name ’SNP_StrandBias’ --filter-name ’SNP_HaplotypeScoreHigh’ --filter-name ’SNP_MQRankSumLow’ -- filter-name ’SNP_ReadPosRankSumLow’ --filter-name ’SNP_HardToValidate’ --filter-name ’SNP_LowCoverage’ -- filter-name ’SNP_VeryLowQual’ --filter-name ’SNP_LowQual’ --filter-name ’SNP_SOR’ -cluster 3 -window 10”.

Annotation of variants was done using Ensembl Variant Effect Predictor (VEP)^37^ (v.112.0) with settings = "--vcf -- plugin AlphaMissense,file --plugin NMD --plugin dbNSFP". The full pipeline is available at https://github.com/ToolsVanBox/ASAP. The version of ASAP used was v.1.0.4.

### Somatic mutation filtering

Somatic mutation calls were generated using SMuRF (v3.0.0) to ensure high-confidence variant identification (available at www.github.com/ToolsVanBox/SmuRF). Variants were retained if they met the following criteria: (A) assigned a GATK Phred-scaled quality score of at least 100, (B) had a mapping quality of ≥60 for 30x coverage data or ≥55 for 15x coverage, (C) had minimum base coverage of 10 (30x) or 5 (15x), (D) exhibited a GATK genotype quality of 99 (for heterozygous calls) or 10 (for homozygous calls) in both the tumour sample and matched normal, and (E) showed no supporting reads in the matched control. Finally, mutations with low variant allele frequencies (VAF) were removed to obtain a set of clonal mutations. The VAF cut-off for both single-cell and bulk samples was 0.15, as mutations below this cut-off could be technical artifacts.

### Mutation filtering for PTA single-cell WGS data

For single cells, our in-house developed pipeline PTATO (v.1.3.3) was applied, which takes the VEP VCF files produced by ASAP (v.1.0.4) and filters these using germline mutations combined with a random forest and walker to separate real somatic mutations from amplification-induced artifacts. For a full description of this pipeline, see our previously published publication^23^.

### Quality control of single-cell WGS data

To ensure high-confidence single-cell genomic profiles, we first excluded samples with a callable loci fraction below 0.45 (n=13). The cutoff was determined based on the distribution of the callable loci fraction among all samples (Supplementary Fig.10a). Next, for the remaining samples, we fitted a logistic regression model to describe the relationship between mean coverage and callable genome fraction and removed samples with residuals below −0.05 as having low mapping quality (n=37) (Supplementary Fig. 10b). Again, the cutoff here was selected based on the distribution (Supplementary Fig. 10c). B-allele frequency (BAF) plots were then manually reviewed and samples with high allelic imbalance were filtered out (n=25). Finally, with the remaining samples we assessed variant-level quality by calculating the total variation distance (TVD) between each sample’s VAF histogram and the cohort median distribution. Samples with a TVD exceeding the median + 3.5 × Median Absolute Deviation (MAD) were flagged as outliers (n=7) (Supplementary Fig. 10d).

### Per sample MYC::IG mutation status

For bulk WGS samples, the location of the *MYC::IGH* translocation was manually examined using IGV (v.2.8.2). Patient-specific primers were designed that flank the translocation breakpoints to assess the presence or absence of the *MYC::IGH* rearrangement in single-cell samples. In addition, the hallmark translocation was manually reviewed and validated using IGV (v.2.8.2)^38^. Samples were classified as translocation-positive if at least one pair of reads spanned the breakpoint region (Supplementary Fig. 11).

### EBV status

To assess Epstein–Barr virus presence and potential genomic integration, we used VIRUSBreakend^16^, a computational tool for detecting viral sequences and integration breakpoints from WGS data. As a positive control, we analysed a previously sequenced bulk WGS sample from the AHH-1 cell line, known to be EBV-transformed (NCBI Taxonomy ID: 10376). As expected, EBV reads were detected in the AHH-1 sample. In contrast, no evidence of EBV DNA or viral integration was found in any of the patient-derived bulk WGS samples analysed in this study.

### B-cell receptor repertoire

BAM files for each single-cell or bulk sample were first down-sampled to the TCR and BCR loci using a custom BED file. The locus-specific BAMs were then converted to paired-end FASTQ files. Receptor reconstruction was performed with MiXCR (v.4.3.2)^39^. For each clone, the top-scoring V, D and J genes, as well as the amino acid CDR3 sequence, were extracted. Clones with CDR3 sequences containing incomplete (“_”) and/or stop (“*”) codons were filtered out from downstream analysis. BCR repertoires were visualized using the R package *ComplexHeatmap* v.2.22.0^40^.

### Phylogenetic tree construction

Phylogenetic reconstruction of normal and malignant single cells from the same patient was performed using CellPhy (v.0.9.254)^41^, which relies on RAxML-NG for maximum likelihood inference. CellPhy considers allelic dropout and amplification errors on a per-sample basis to estimate the most likely tree structure. Analyses were performed using phred-scaled genotype likelihoods (PL) under the default “GT16 + FO + E” model. For each tree, 100 bootstrap iterations were generated, and the proportion of replicates supporting each split is shown in Supplementary Fig. 12, providing a measure of node confidence. Since CellPhy assigns mutations based upon where it will provide the most information on which split occurs and not based on where the mutations actually are within the tree, we addressed this by using our own CellPhyWrapper function available at https://github.com/ToolsVanBox/CellPhyWrapper.

Trees were rooted using the “root” function from the *treeio* R package, with MSCs used as the outgroup. To validate tree topology, variant allele frequencies (VAFs) from bulk tumour samples were assessed. Clonal (trunk) branch mutations displayed variant allele frequencies (VAFs) of approximately 0.5, consistent with their presence in all tumour cells. By contrast, mutations assigned to intermediate branches - positioned between trunk and private branches - were either undetectable or detected at low VAFs in bulk sequencing, reflecting the resolution limits of this method. Private branch mutations were almost uniformly undetectable in bulk, indicating their restriction to individual malignant cells (Supplementary Fig. 3).

To account for differences in sequencing sensitivity between cells, we corrected branch lengths using callable loci information. Specifically, we calculated a node-specific detection power, 𝑝, as 𝑝 = 1 − Π(1 − 𝑠_𝑖_), where 𝑠_𝑖_ is the callable-loci fraction of sample 𝑖, and scaled the original CellPhy branch lengths by this value to yield callable- normalised branch lengths.

### SBS and INDEL driver annotation

Firstly, single base substitution (SBS) and insertion–deletion (INDEL) variants were retained if the Variant Effect Predictor (VEP) assigned an impact of *Moderate*, *High*, or *Likely pathogenic*. Then, somatic variants absent in matched germline controls and passing standard quality filters were kept. To focus on bona fide drivers, we filtered this set on known driver genes. Specifically, we filtered against a panel of 86 Burkitt lymphoma driver genes curated from three datasets: IntOGen, Nicole, et al. ^7^ and Panea, et al^11^ (Supplementary Table 3). Finally, only protein-coding, non-synonymous variants - including missense, nonsense, frameshift, start/stop lost, and inframe INDELs - were retained for downstream analysis, while synonymous and noncoding variants were excluded. Oncoplots were generated using the *maftools* package^42^ (v2.22.0). For single-cell WGS data, variants were additionally required to be absent from all patient-matched normal B cells and present in at least one malignant cell. In addition, SBS and INDEL driver events were manually inspected in IGV (v.2.8.2) to confirm their position within the phylogenetic tree. If a driver mutation was not detected in a given tumour cell despite being present in other cells within the same clade, we attributed this absence to technical limitations, following the principle of parsimony (Occam’s razor) based on the phylogenetic reconstruction. All SBS and INDEL driver mutations identified in the bulk and single-cell samples are provided in Supplementary Tables 2 and 3, respectively.

### CNV annotation

For CNVS, we aggregated PURPLE-derived^43^ CNV segment outputs per sample, kept only autosomal chromosomes and iteratively merged adjacent segments (≤30 Mb gaps; ≥10 kb length). Shared CNV regions were defined within patient groups by building overlap graphs of segments that reciprocally overlapped ≥80% and extracting connected components. Regions were then mapped to cytobands. Lastly, copy-number states were harmonized to gain (red), loss (blue), or loss of heterozygosity (LOH) (green), collapsed per (cytoband, sample), and visualized as cohort-specific heatmaps (Supplementary Fig. 13). All CNVs identified in the bulk and single-cell samples are provided in Supplementary Tables 2 and 3, respectively. As for SBS and INDEL drivers, we followed the principle of parsimony (Occam’s razor) based on the phylogenetic reconstruction to assign CNVs in the branches.

### Selective pressure analysis

To assess selective pressures on coding mutations in our bulk WGS data, we applied the *dNdScv* R package (v0.0.1.0)^20^ using the hg38 reference genome. Here, only autosomal SBS variants were included for standard dN/dS analysis. The analysis was performed against the hg38 reference genome using default trinucleotide context corrections. For each gene, the ratio of nonsynonymous to synonymous substitution rates (dN/dS) was estimated separately for missense and truncating classes. Gene-level selection coefficients were extracted and genes with a global q-value < 0.1 were considered under positive selection. Effect sizes (w) and q-values were log-transformed for visualization, and the most significant class per gene (missense or truncating) was used to construct a volcano plot highlighting genes under positive selection.

To assess selective pressures across tumour evolution, we estimated dN/dS ratios within single-cell WGS– derived phylogenetic branches. Mutations were assigned to phylogenetic branches and grouped into four categories per patient: Normal B-cells, BL Pre-expansion (trunk), BL-Intermediate, and BL-Private. Immunoglobulin loci (IGH, IGK, IGL) were excluded to avoid bias from somatic hypermutation. *dNdScv* was run separately for each group to compute overall dN/dS values and 95% confidence intervals, which were visualized as dot and forest plots.

### SBS burden analysis

We estimated per-cell mutational load from PTATO-filtered VCFs by retaining autosomal variants and correcting for each cell’s callable-genome fraction, thus yielding SBS per callable genome. To place tumour loads in context, we combined our normal and malignant B-cells with healthy naïve/memory B-cell colonies from Machado et al.^13^ and with bulk BL tumours. For bulk tumour samples, VAFs were modelled per sample with a Gaussian mixture to identify the peak with the highest mean (i.e. clonal peak) and defined a tolerance band (3 x SD of peak) to determine clonal variants within that peak (Supplementary Fig. 14). Age effects were modelled using linear mixed-effects regression with the naïve B-cells from healthy donors fit provided the age-specific baseline. Then, per sample, we computed the Observed/Expected (O/E) ratio (i.e. mutation load/baseline at that age). Group differences were assessed with two-sided Wilcoxon tests.

### Relative timing of CNVs in bulk samples

The relative timing of CNVs found in bulk WGS samples was inferred using the *MutationTimeR* package (v1.0.2)^44^. Timing was estimated from the proportion of mutations in different copy-number states within each CNV, which reflects the chronological order of mutational acquisition under the assumption of a constant mutation rate. Copy-number profiles generated by PURPLE were smoothed by merging adjacent segments with copy-number ratios between 0.9 and 1.1, rounding values to integers, and combining neighbouring events with equal CN. Small segments (<1 Mb) were subsequently merged with the nearest larger event to further reduce noise. The resulting list of CNVs was used as input to generate a circos plot depicting the frequency of CNVs in the bulk BL samples. MutationTimeR was then applied to all samples using the processed CN events and previously identified mutations. Low-confidence events with confidence intervals >0.8 and widths <7 Mb were removed (Supplementary Table 1).

### Mutational signature extraction and refitting

Mutational signature analysis was carried out using the *MutationalPatterns* (v.3.16.0) R package^45^. 96- trinucleotide profiles were extracted from the SBS mutations in the single-cell WGS samples to identify the landscape of mutational contexts. To increase our confidence in extracting mutational signatures, we included additional mutational profiles: bulk WGS of Burkitt lymphoma samples from the Princess Máxima Center for Pediatric Oncology cohort (n=21), a collection of previously published healthy T-cell, B-cell and HSC reference samples (n=734)^13^ and genomes from 7 blood cancer types, including Burkitt lymphoma, follicular lymphoma, diffuse large B cell lymphoma, chronic lymphocytic leukaemia (mutated), chronic lymphocytic leukaemia (unmutated), and acute myeloid leukaemia (AML) and multiple myeloma (MM) (n=223)^46,47^.

*De novo* mutational signature extraction using non-negative matrix factorization (NMF) was performed to identify the signatures that are present in the samples. Extraction was done by applying the “extract_signatures” function with options “rank = 8, nrun = 100. Cosine similarities between the extracted signatures and COSMIC SBS signatures (version 3) were calculated to identify previously described signatures among the extracted signatures. Signature SBSblood was also included^13^. From this, the following signatures were identified: SBS1- like (cosine 0.9), SBS9-like (cosine 0.86), SBS18-like (cosine 0.9), SBSblood (cosine 0.92), SBS17b-like (cosine 0.91), SBSA, SBSB, and SBSC. The SBSC profile showed high similarity with SBS2 (cosine 0.85) and SBS7a (cosine 0.83) and was therefore substituted for these two signatures (Supplementary Fig. 15a). SBSA and SBSB had low cosine similarity to any known signature. Together, the following set of signatures was used for signature refitting: SBSA, SBSB, SBS1, SBS2, SBS7a, SBS9, SBS17b, SBS18 and SBSblood.

To determine the contribution of these signatures to the single-cell and bulk WGS Burkitt samples, strict bootstrapped refitting was performed using 100 bootstraps per sample with “max_delta = 0.002”. SBSA, SBSB and SBS2 all had less that 5% in each genome and were excluded from the analysis and the signature proportions were re-estimated using the remaining 6 signatures: SBS1, SBS7a, SBS9, SBS17b, SBS18 and SBSblood. The cosine similarity between the original and reconstructed matrices was >0.8 in all samples (mean: 0.94; range: 0.82-0.98) (Supplementary Fig. 15b). To test signature robustness, we added a non-extracted signature (including the artefact signatures due to PTA-based whole genome amplification) one-at-a-time and recalculated the cosine similarity, as well as removing one of the six extracted signatures individually. For each sample, we quantified the Δ cosine similarity relative to the original refitting using the 6 extracted signatures and visualized the per-sample changes. Indeed, adding the artefact signatures didn’t improve the refitting, while removing any of the 6 signatures did worsen the cosine similarity (Supplementary Fig. 16).

### Mutational signature contribution within phylogenetic trees

Fitting mutational signatures to the mutations of single branches of the tree was done with the *MutationalPattern*s^45^ function “fit_to_signatures_strict_tree” with option “max_delta = 0.002”. For this analysis, branches were assigned to one of four categories: “Normal B-cell”, “Pre-BL”, “BL-Early”, or “BL-Late”. Normal B-cell branches comprised all private branches derived from normal B-cells. The BL-Trunk category corresponded to the ancestral branch preceding the MRCA of the tumour. BL-Early branches were defined as those arising after the MRCA but prior to the emergence of private branches. BL-Late branches represented private branches restricted to individual Burkitt lymphoma cells. To contrast normal subgroups with BL compartments, we built per-patient signature matrices for BL trunk, intermediate, and private branches, converted them to within-group proportions, and merged them with normal (SBS9^+/-^). For each signature, we then fitted a beta-GLMM with Group as a fixed effect and Patient as a random intercept and derived estimated marginal means. To test if changes in relative contribution between normal/malignant groups were significant a two-sided Wald z-test was used.

### Timing branches

To estimate the timing of mutational events, we reconstructed ‘time-based’ phylogenetic trees using the *rtreefit* R package (https://github.com/nangalialab/rtreefit). This approach converts phylogenetic trees with branch lengths in molecular time (number of SBS mutations) into trees scaled in chronological time (years). In brief, the method simultaneously estimates wild-type and mutant mutation rates (the number of SBSs accumulated per year) and absolute branch durations within a Bayesian, per-individual tree model. It assumes that the number of mutations assigned to each branch follows a Poisson distribution while constraining the total root-to-tip length to match the patient’s age at sampling. This was calculated as the patient age plus 9 months, to account for the fertilization age.

We defined the branch absolute mutation lengths as the sum of SBS1 and SBSblood contributions, representing linear, clock-like mutational processes, and the age at sampling as being post-conception. Furthermore, we accounted for an elevated mutation rate during embryogenesis by assuming an excess mutation rate through development. Finally, the mutant clade was defined as not including the tumour trunk, so the method assumed that the mutation rate switch from normal to mutant occurred at the end of the trunk. For donors with two anatomically distinct tumour sites (PRN4 and P856), rate switches were introduced at the end of the shared branches corresponding to each site. No switch was placed for the diagnostic samples of P856 and PIA9, as tumour cells had a similar mutation burden compared to healthy B-cells.

We noted that for the larger trees of patients PJBU, PVA9 and PIA9, the algorithm took a long time when all cells were included, but that time dropped drastically (from ∼2 months to ∼2 minutes) when one cell was removed. Therefore, we removed normal B-cells PIA9GTDBBC56, PJBUGTDBBC73 and PVA9GTDBBC74 for PIA9, PJBU and PVA9, respectively. To test whether removing a normal cell from a tree does not influence timing estimates, we tested this on P3G6 and found no difference in MRCA timing (Supplementary Fig. 17). The *rtreefit* algorithm was run with 12 chains and 20,000 iterations per chain.

## Data availability

Data will be made available through the EGA (accession number pending).

## Code availability

Code created for analyses are available at GitHub (https://github.com/ProjectsVanBox/Burkitt_scWGS).

## Acknowledgments

This work was funded by the Foundation Kids Cancer Free (KiKa; no. 424) to Ruben van Boxtel and Friederike Meyer-Wentrup. In addition, this work was supported by Stichting Maas and the Oncode institute. Ruben van Boxtel is a New York Stem Cell Foundation – Robertson Investigator. This research was supported by The New York Stem Cell Foundation.

## Author contributions

A.S.S., F. M. and R.B. conceived and initiated the project. A.S.S and R.B. designed the experiments. A.S.S., L.T., M.V. performed the experimental work. A.S.S., M.J.R., R.H., J.K.K., S.J. performed the data analysis. F.M. provided clinical data. F.M. and R.B supervised the project. A.S.S., F.M. and R.B. drafted the manuscript.

## Ethics declarations

Ruben van Boxtel is a scientific advisor to BioSkryb Genomics and Hartwig Medical Foundation.

---

**Supplementary Figure 1.**
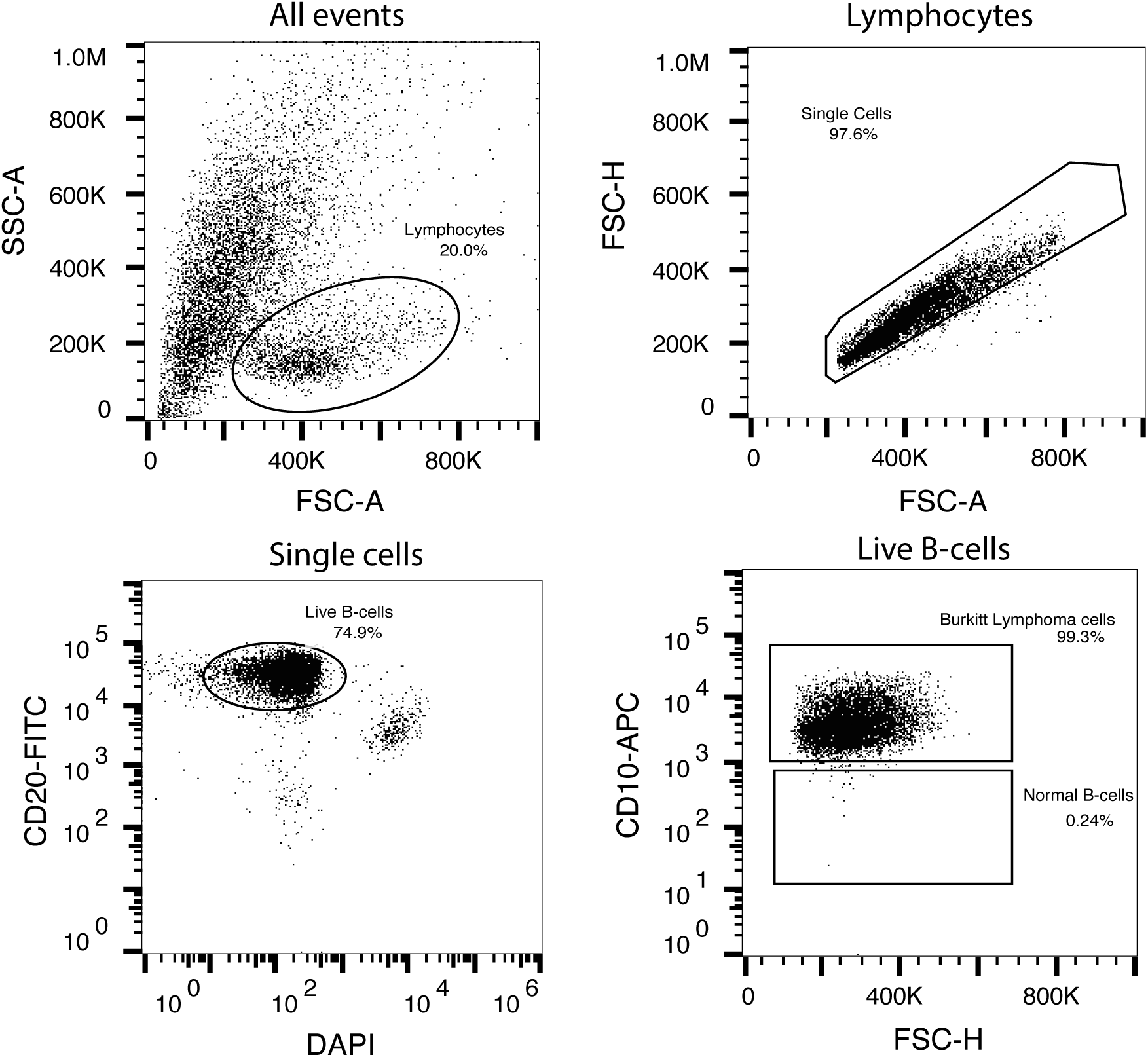
Example of gating strategy for flow-sorting both normal B-cells and Burkitt lymphoma (BL) cells from patient PJBU. First, lymphocytes were selected based on side scatter area (SSC-A) and forward scatter area (FSC-A). Then, singlets were selected based on forward scatter heigth (FSC-H) and forward scatter area (FSC-A). DAPI negative and CD20-FITC positive cells were gated to select for live B-cells. Finally, CD10-APC was used to enrich for BL (CD10 positive) and normal B-cells (CD10 negative).

**Supplementary Figure 2.**
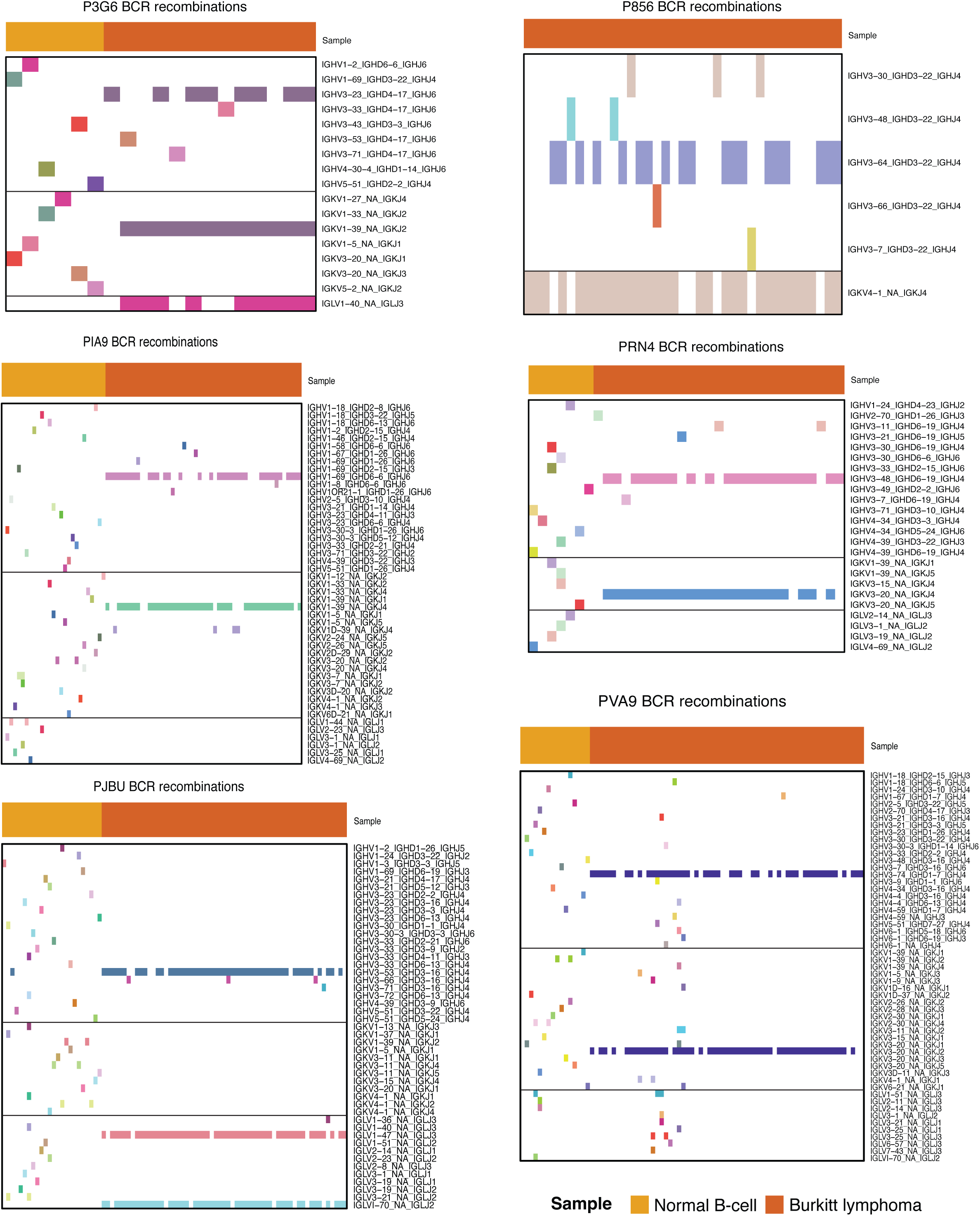
B-cell receptor (BCR) rearrangements, including the immunoglobulin heavy and lambda/kappa light chain rearragnements, present in each normal B-cell and Burkitt lymphoma (BL) cell.

**Supplementary Figure 3.**
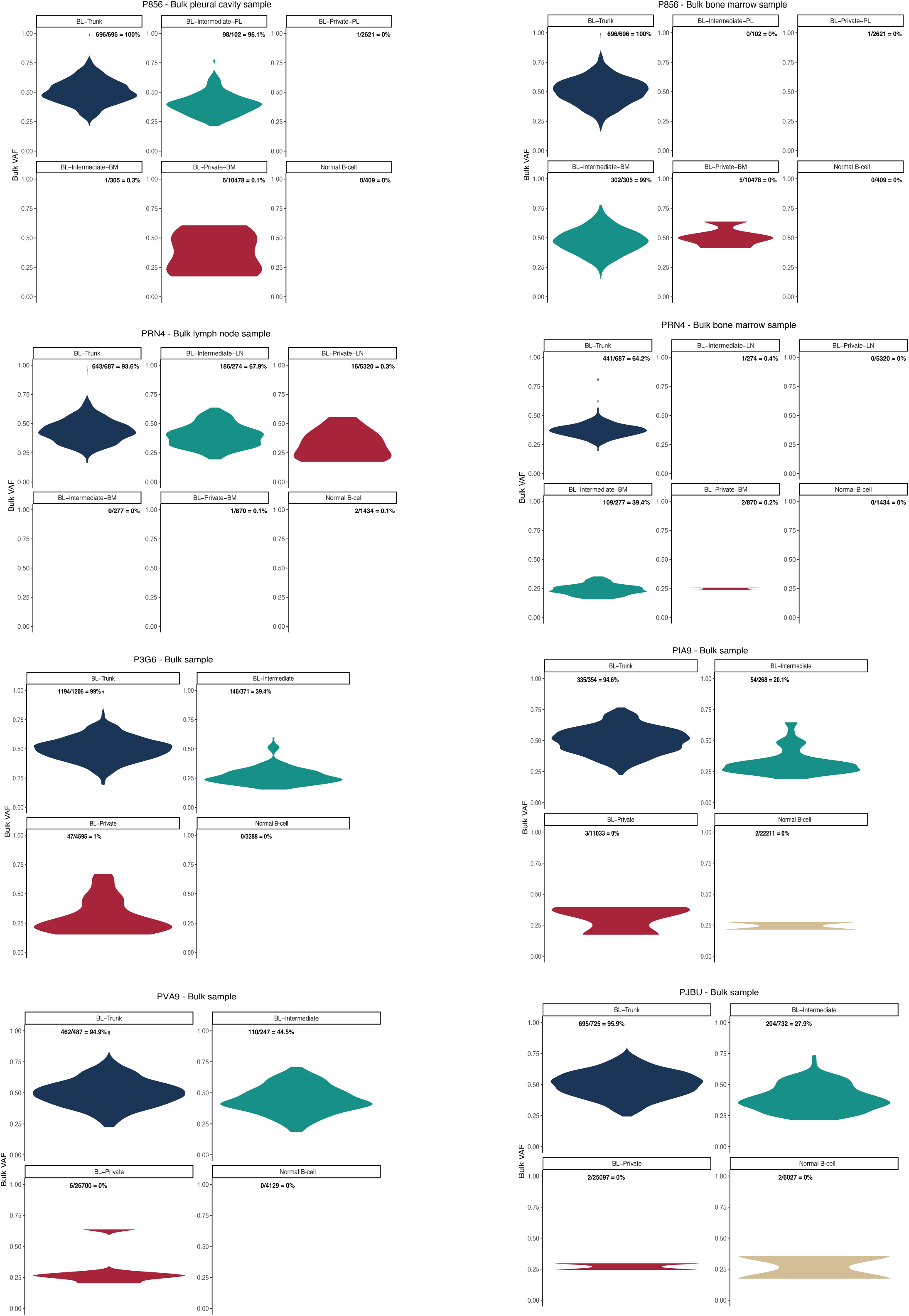
Variant allele frequency (VAF) of branch mutations in the corresponding bulk tumour sample for each patient. For patients P856 and PRN4, two bulk tumour samples were available due to multi-site involvement. Branch mutations from phylogenetic trees were divided into Burkitt lymphoma (BL) trunk mutations, BL intermediate mutations, BL private mutations and normal B-cell private mutations.

**Supplementary Figure 4.**
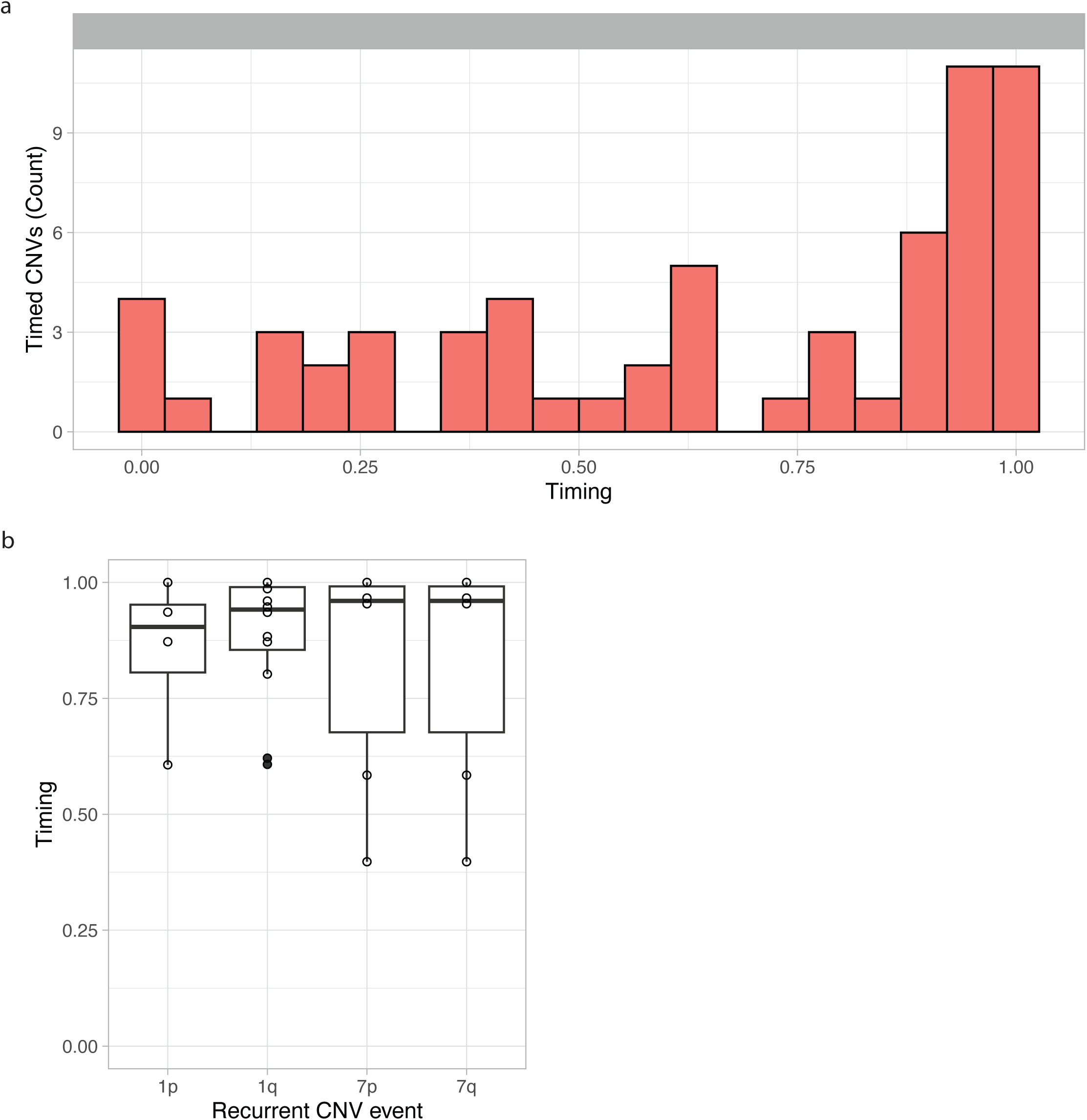
**a**, Relative timing of copy number variants (CNVs) based on *MutationalTimeR* analysis. A value of 0 represents the earliest possible occurrence, while a value of 1 represents the latest. **b**, Relative timing of recurrent CNV events in our Burkitt lymphoma cohort.

**Supplementary Figure 5.**
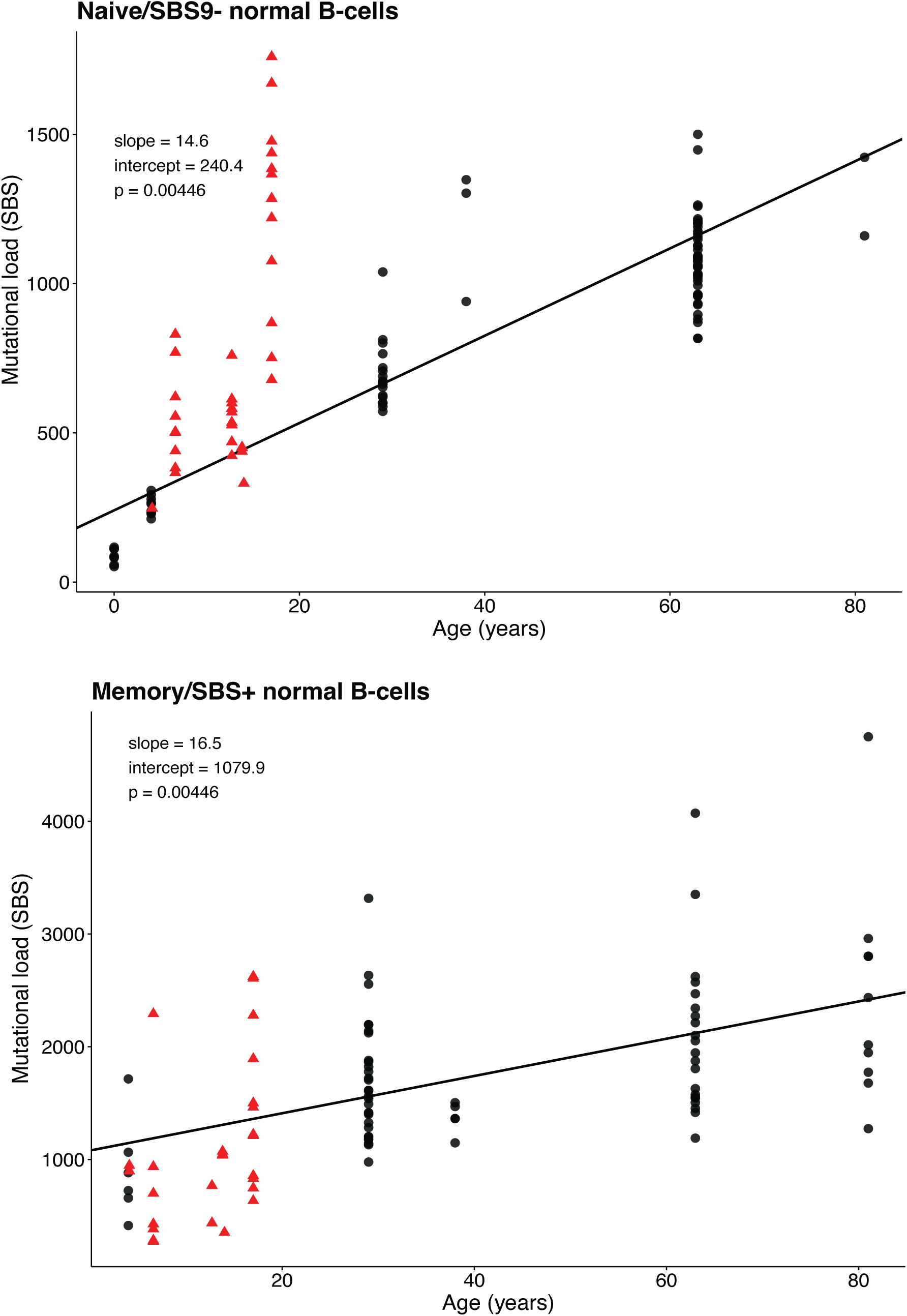
SBS mutational burden per genome. Naive and Memory B-cells were obtained from Machado et. al. (Nature, 2022) and are depicted in black circles. The lines show the fit for naive B-cells (top) and memory B-cells (bottom) using linear mixed-effects models. Normal B-cells from Burkitt lymphoma patients were divided into SBS9- (top) or SBS9+ (bottom) according to their SBS9 mutational process contribution and are depicted in red triangles.

**Supplementary Figure 6.**
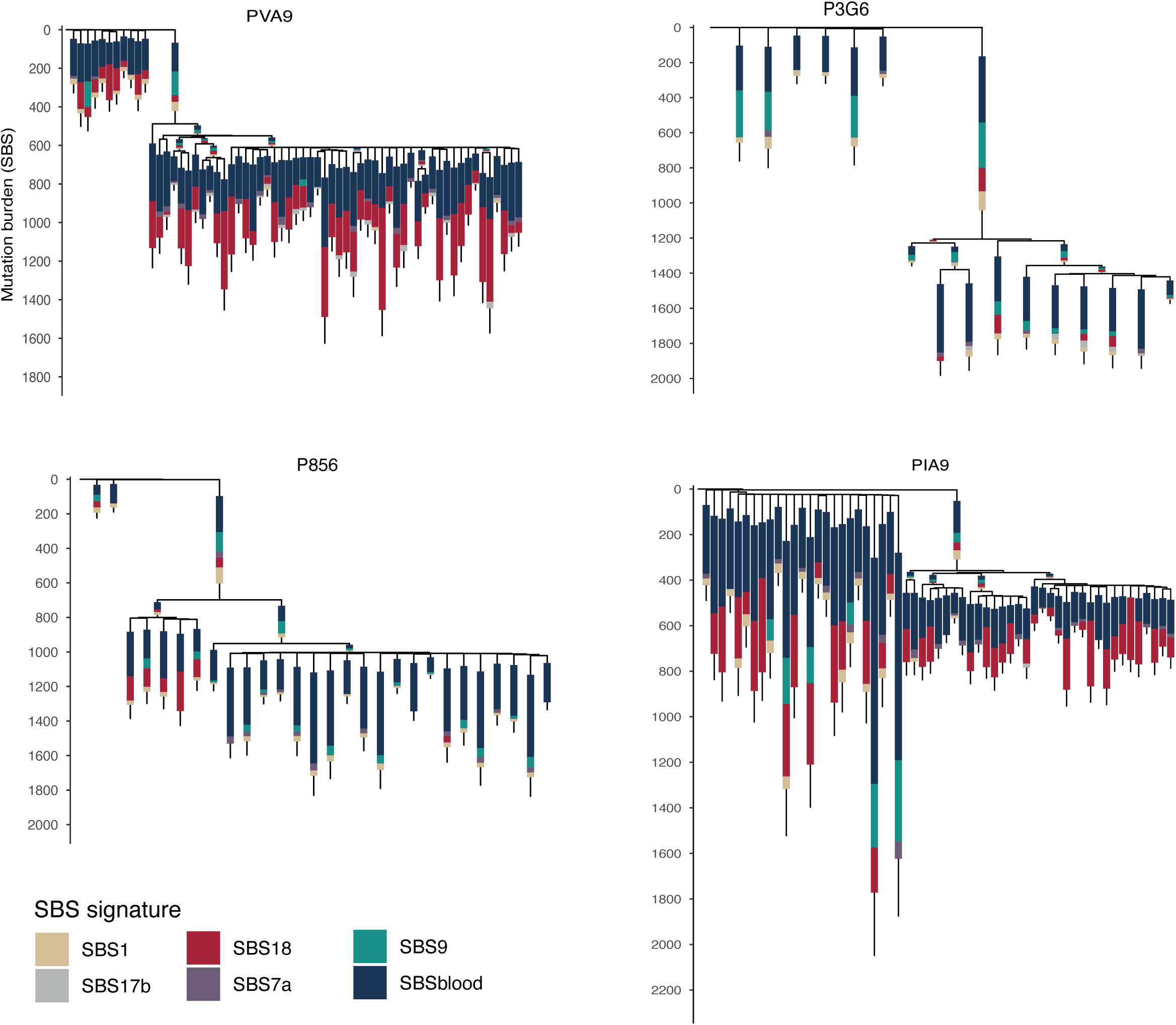
Remaining phylogenetic trees with SBS signature contributions per branch.

**Supplementary Figure 7.**
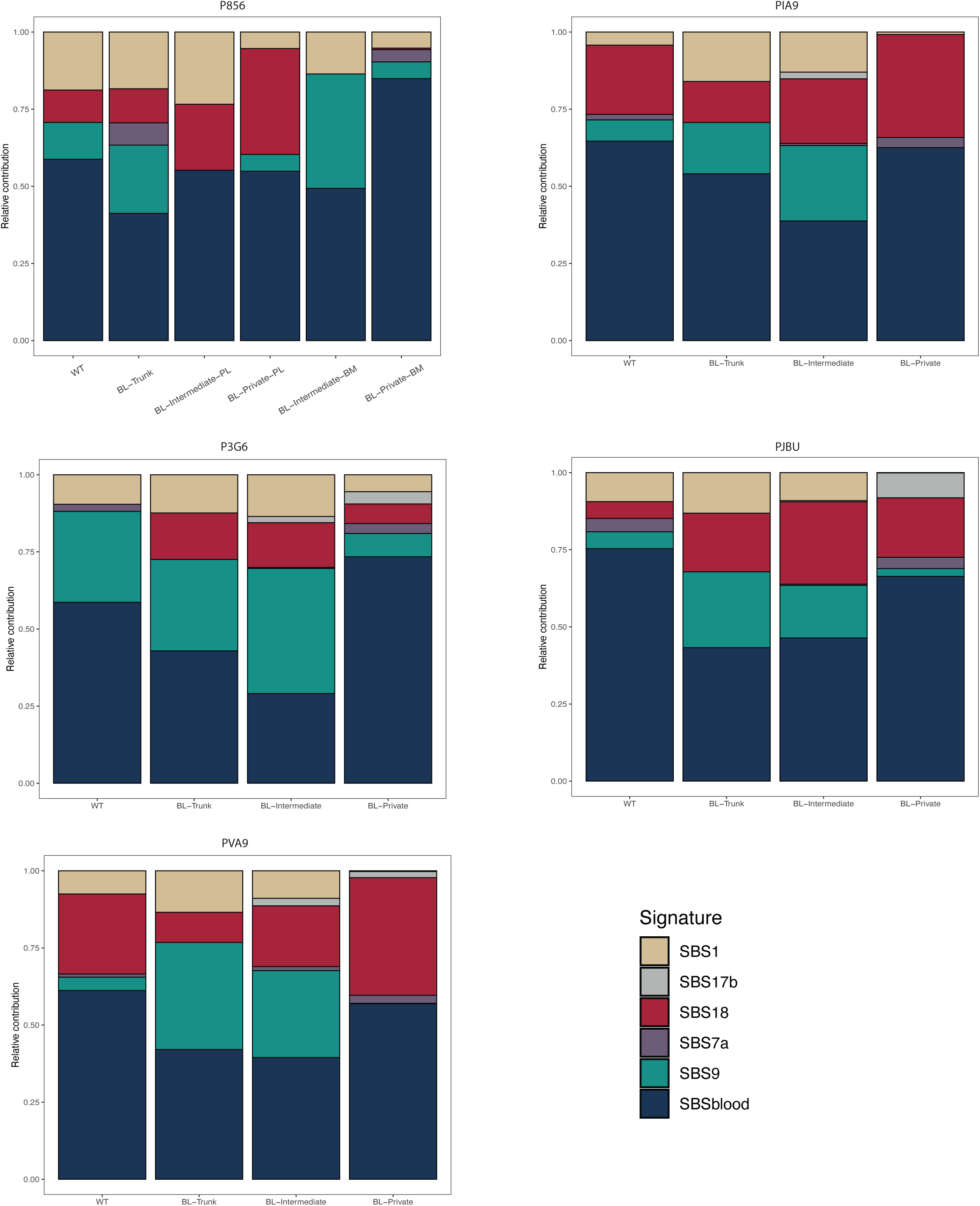
Remaining relative SBS signature contributions per branch group.

**Supplementary Figure 8.**
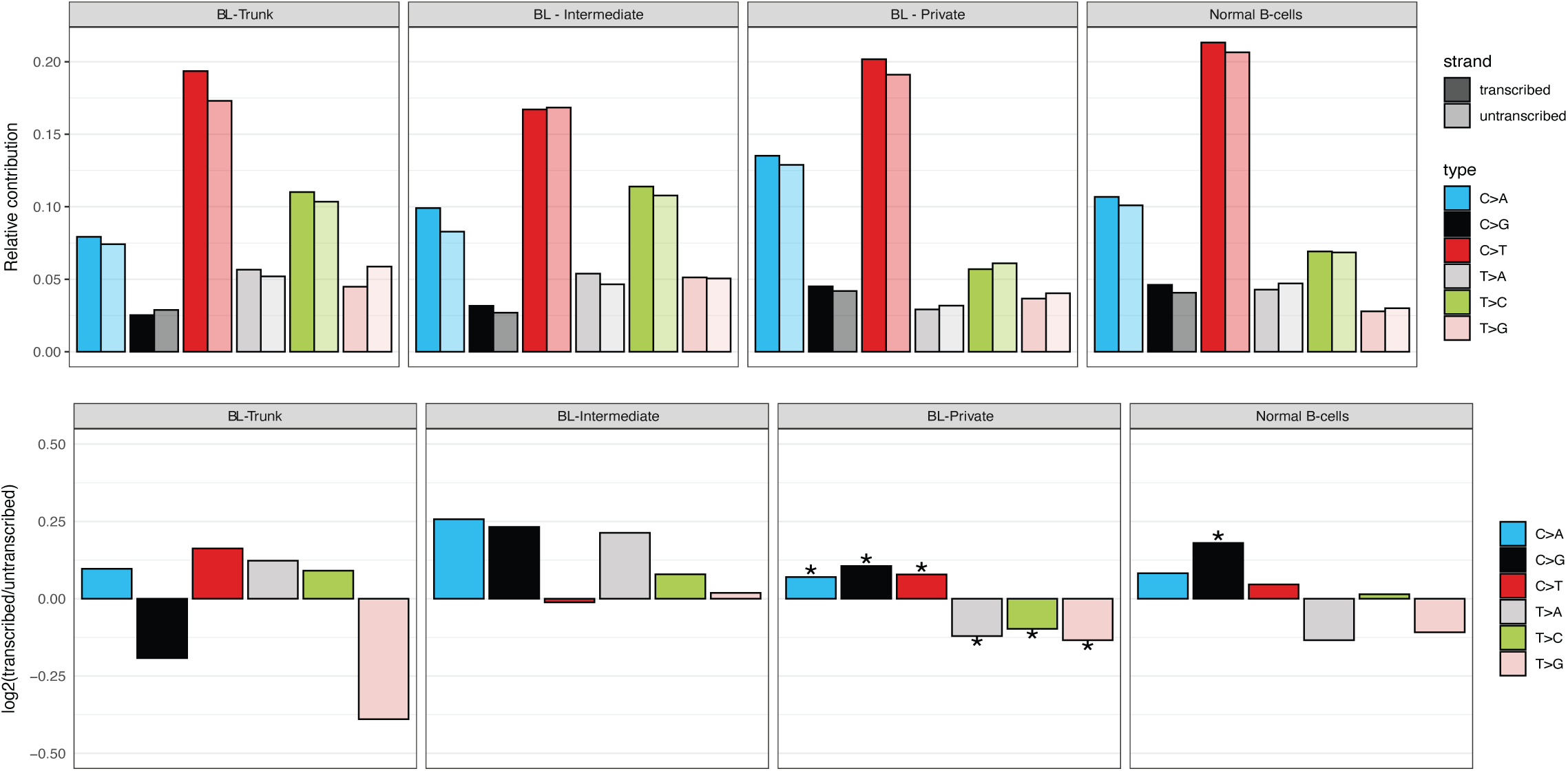
Transcriptional strand bias analysis using *MutationalPatterns*, on distinct branch groups. Top: mutation spectrum with strand distinction. Bottom: effect size (log2(untranscribed/transcribed) of the strand bias. Asteriks indicate significant strand bias.

**Supplementary Figure 9.**
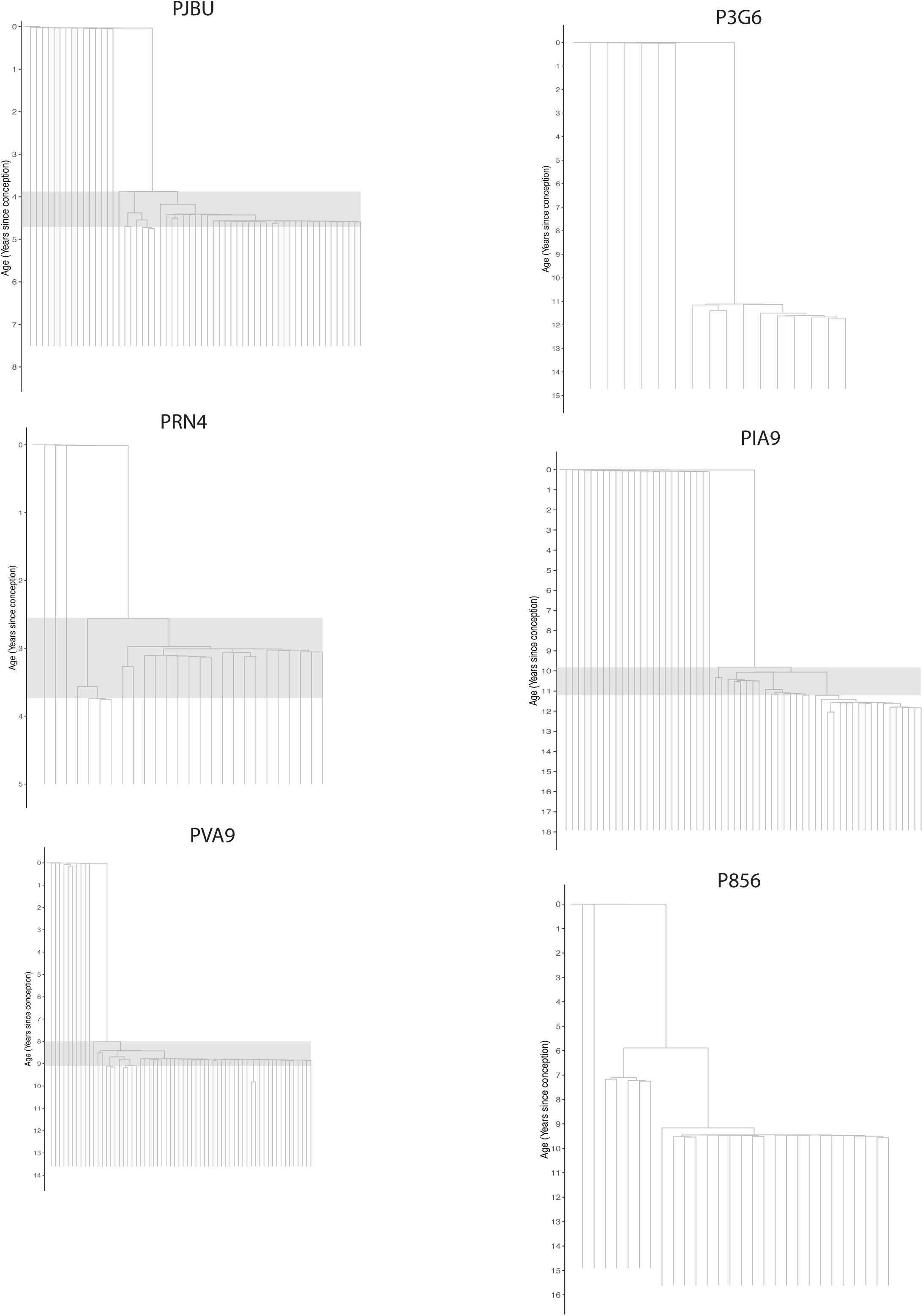
‘Time-based’ trees of all patients, in years post-conception. Grey boxes depict the time window in each subclonal drivers accumulated in a subset of patients.

**Supplementary Figure 10.**
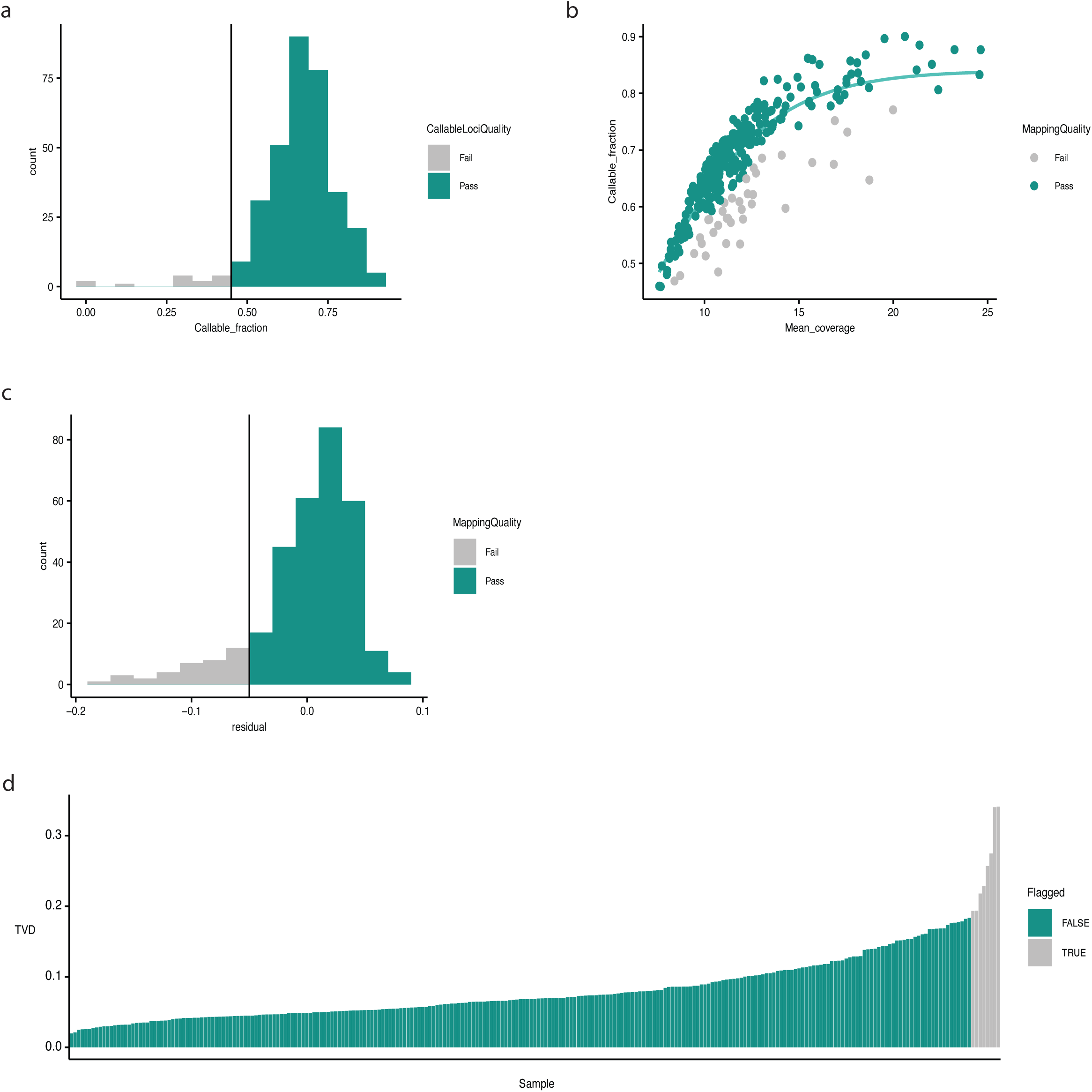
**a**, Histogram showing the distribution of the callable loci fraction across 332 single cells whole genome amplified using Primary Template-directed Amplification (PTA). Cut-off determined was 0.45. **b**, Logistic regression model fitted on mean coverage and callable fraction. Residual is calculated as the difference between the observed callable genome fraction for a sample and the value predicted by the logistic regression model based on its mean sequencing coverage. **c**, Histogram showing the distribution of the residual score for each cell. Cut-off determined was -0.05. **d**, Distribution of the total variation distance (TVD) between each sample’s VAF histogram and the cohort mean VAF distribution. Samples with TVD > median + 3.5 x Median Absolute Deviation (MAD) were flagged.

**Supplementary Figure 11.**
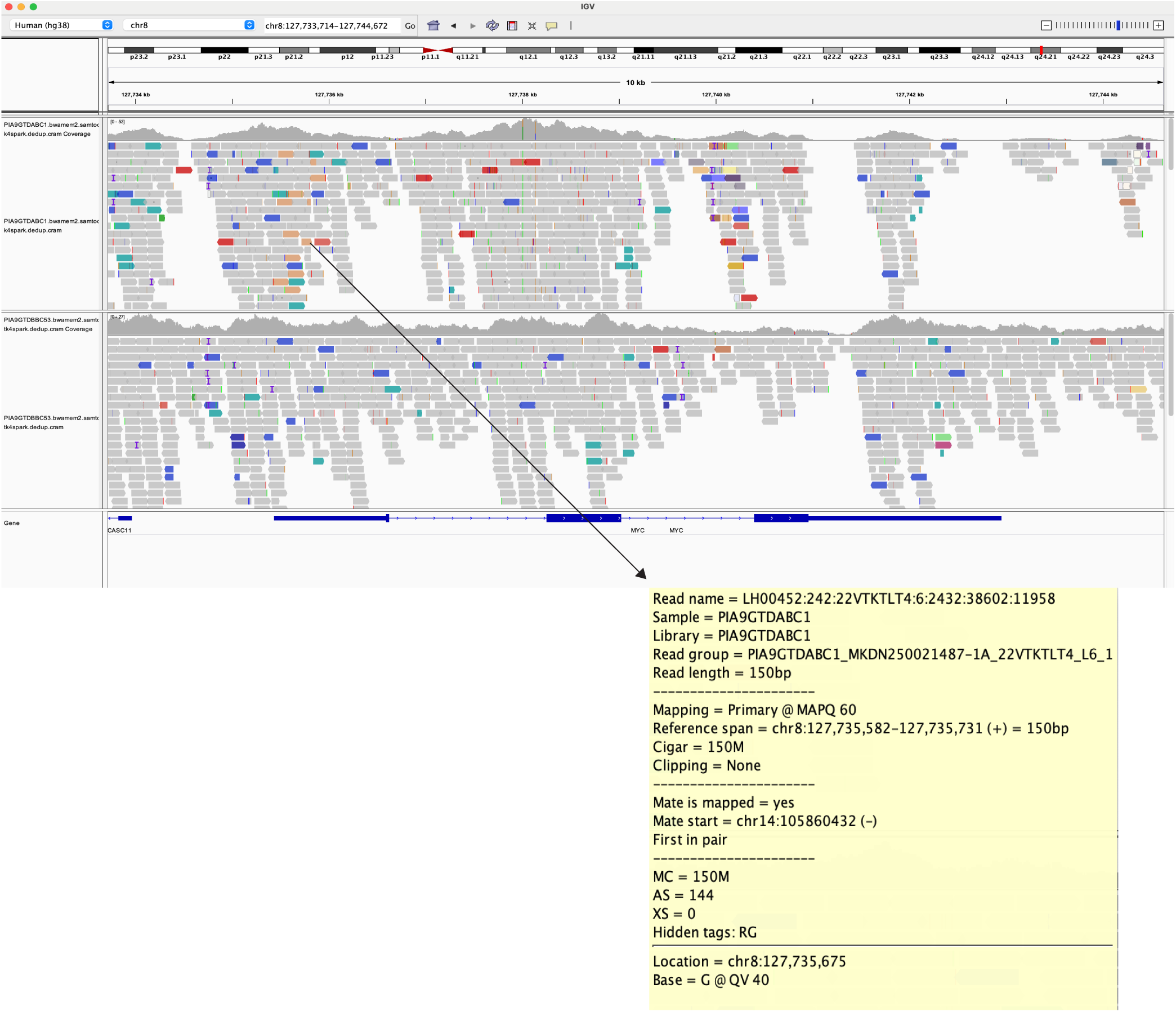
Example of IGV output to determine the presence/absence of the *MYC::IGH* translocation. Top sample (PIA9GTDABC1) is a *MYC::IGH*-positive cell while the bottom sample (PIA9GTDBBC53) is a *MYC::IGH*-negative cell. Reads annotated in orange depict reads that map partly to the *MYC* locus on chromosome 8 and partly to the *IGH* locus on chromosome 14.

**Supplementary Figure 12.**
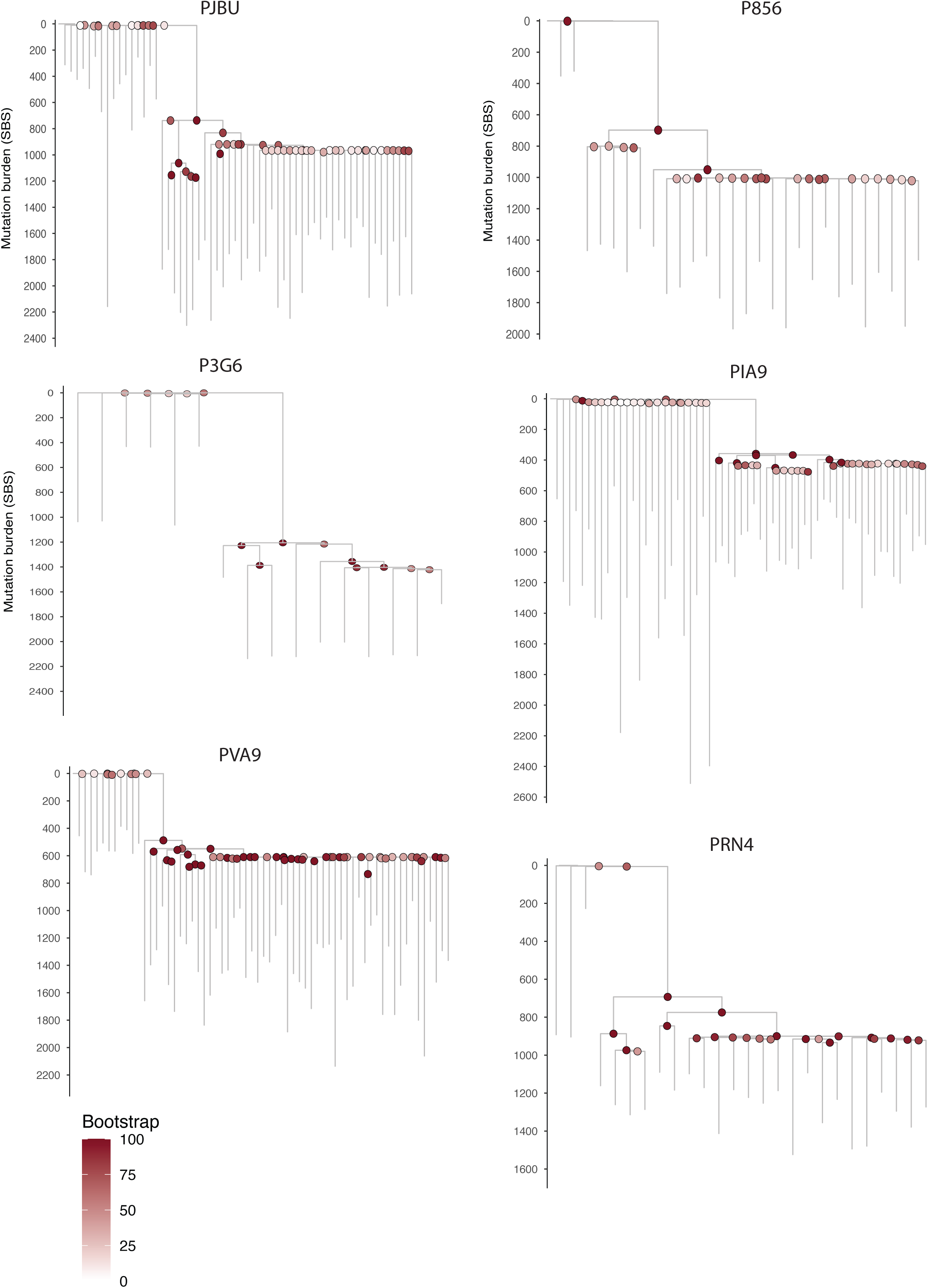
Phylogenetic trees with bootstrap confidence on each node. For each tree, 100 bootstrap iterations were generated.

**Supplementary Figure 13.**
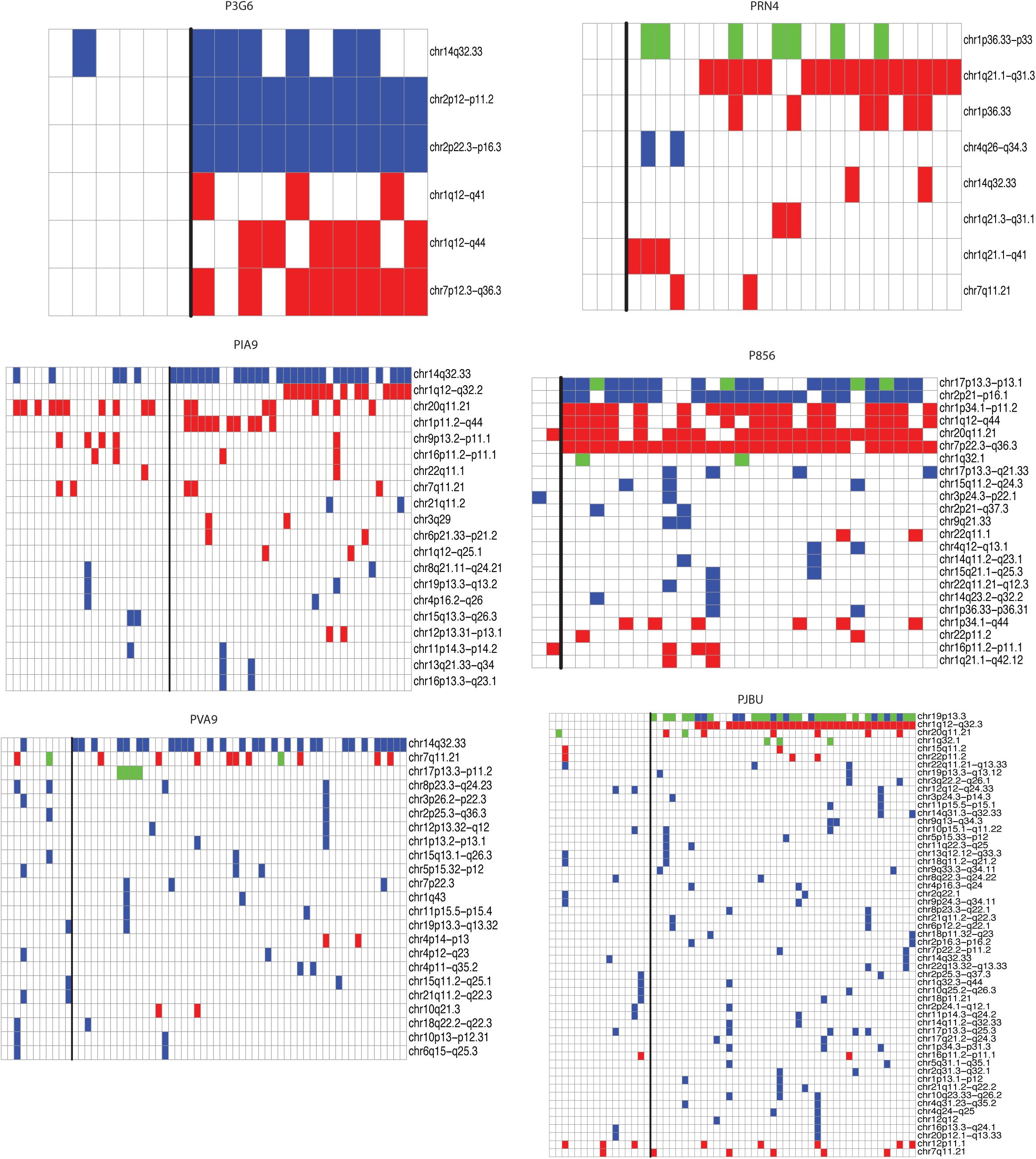
Heatmap showing the presence/absence of a copy number variation (CNV), per cell. CNVs include gains (red), losses (blue), and loss of heterozygosity (green). The black vertical line splits the heatmap between normal B-cells on the left and Burkitt lymphoma (BL) cells on the right. The order of both normal and BL cells corresponds to the order of those cells on the phylogenetic trees.

**Supplementary Figure 14.**
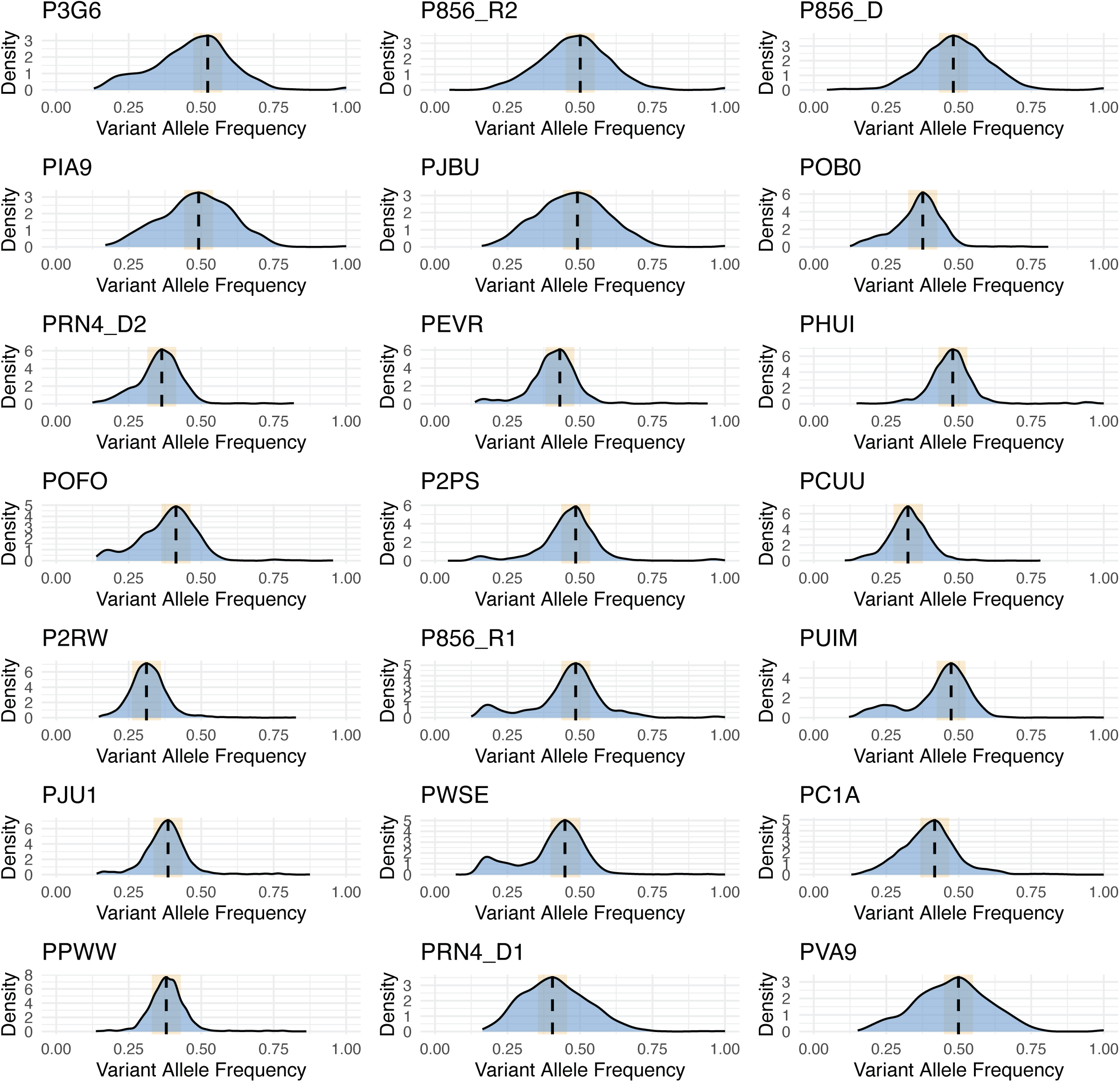
Selection of clonal mutations from bulk WGS daa. ta. VAFs were modelled per sample with a Gaussian mixture to identify the peak with the highest mean (i.e. clonal peak; black dotted line) and defined a tolerance band (3 x SD of peak) to determine clonal variants within that peak (yellow shaded area).

**Supplementary Figure 15.**
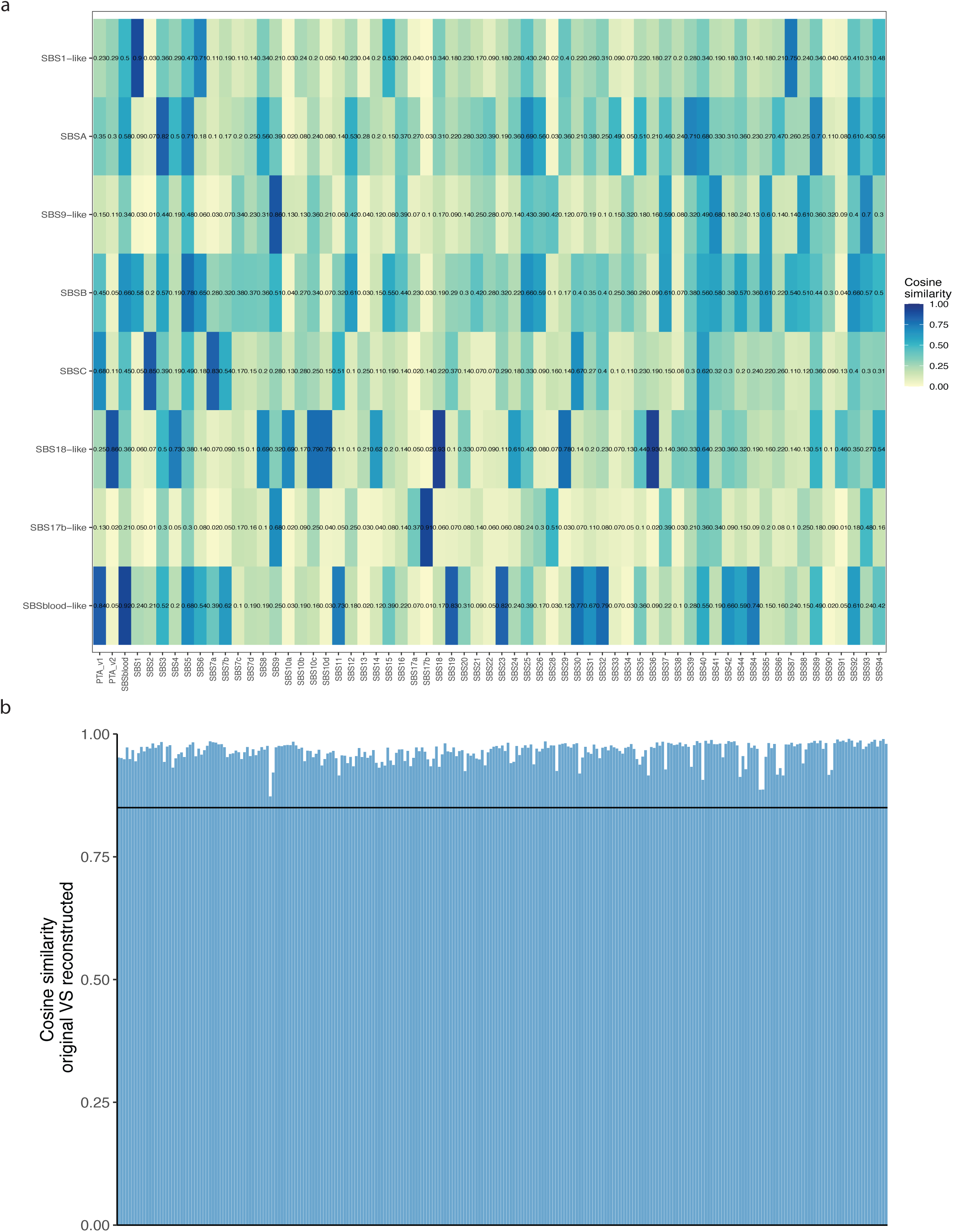
**a**, Cosine similarities between the *de novo* extracted signatures and COSMIC SBS signatures (version 3), including signature SBSblood. **b**, The cosine similarity between the original and reconstructed matrices using the following signatures: SBS1, SBS7a, SBS9, SBS17b, SBS18 and SBSblood. Vertical line depicts the 0.85 threshold.

**Supplementary Figure 16.**
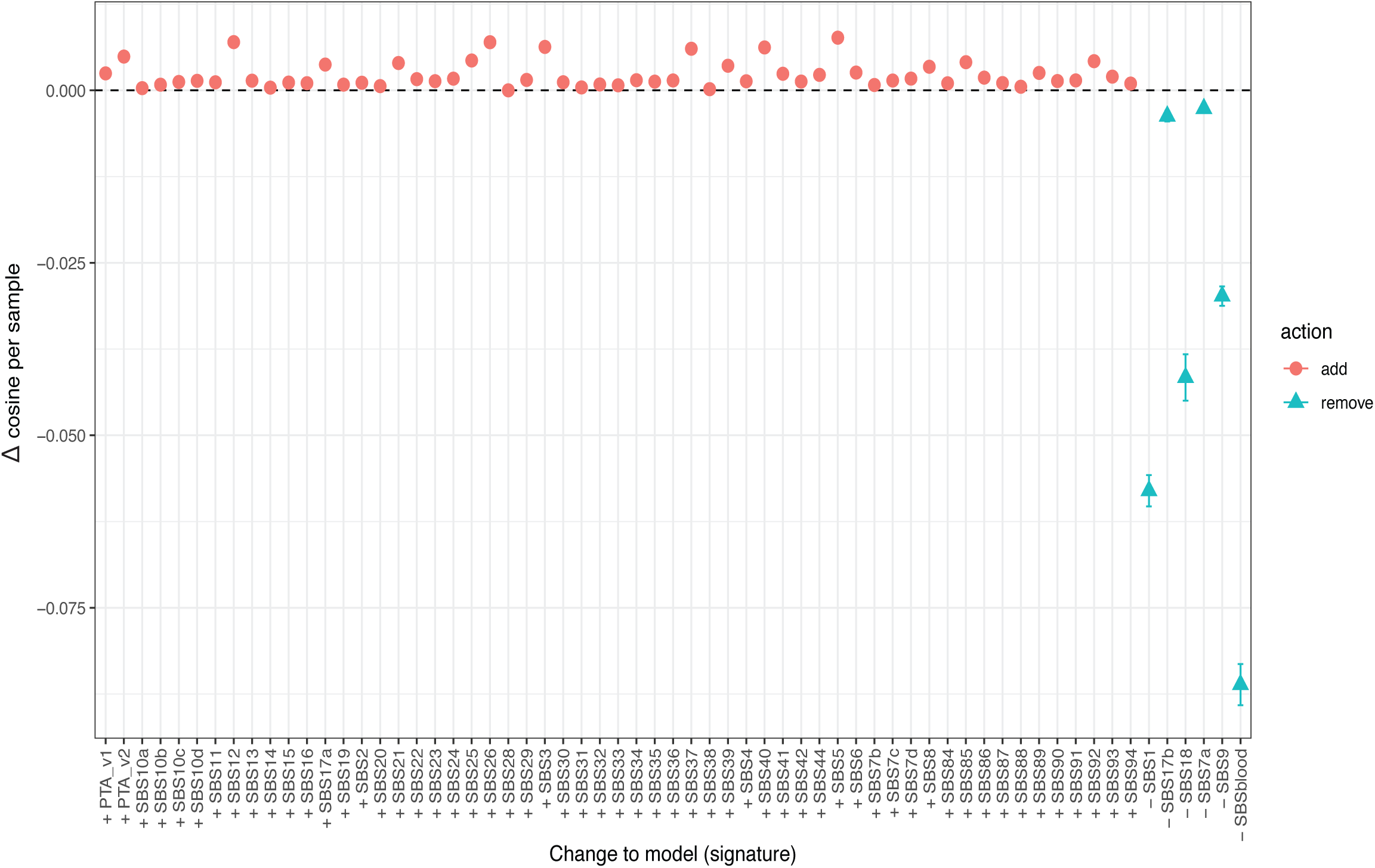
Δ cosine similarity relative to the original refitting. This was done by first adding a non-extracted signature (including the artefact signatures due to PTA-based whole genome amplification) one-at-a-time and recalculating the cosine similarity (red), as well as removing one of the six extracted signatures individually (blue).

**Supplementary Figure 17.**
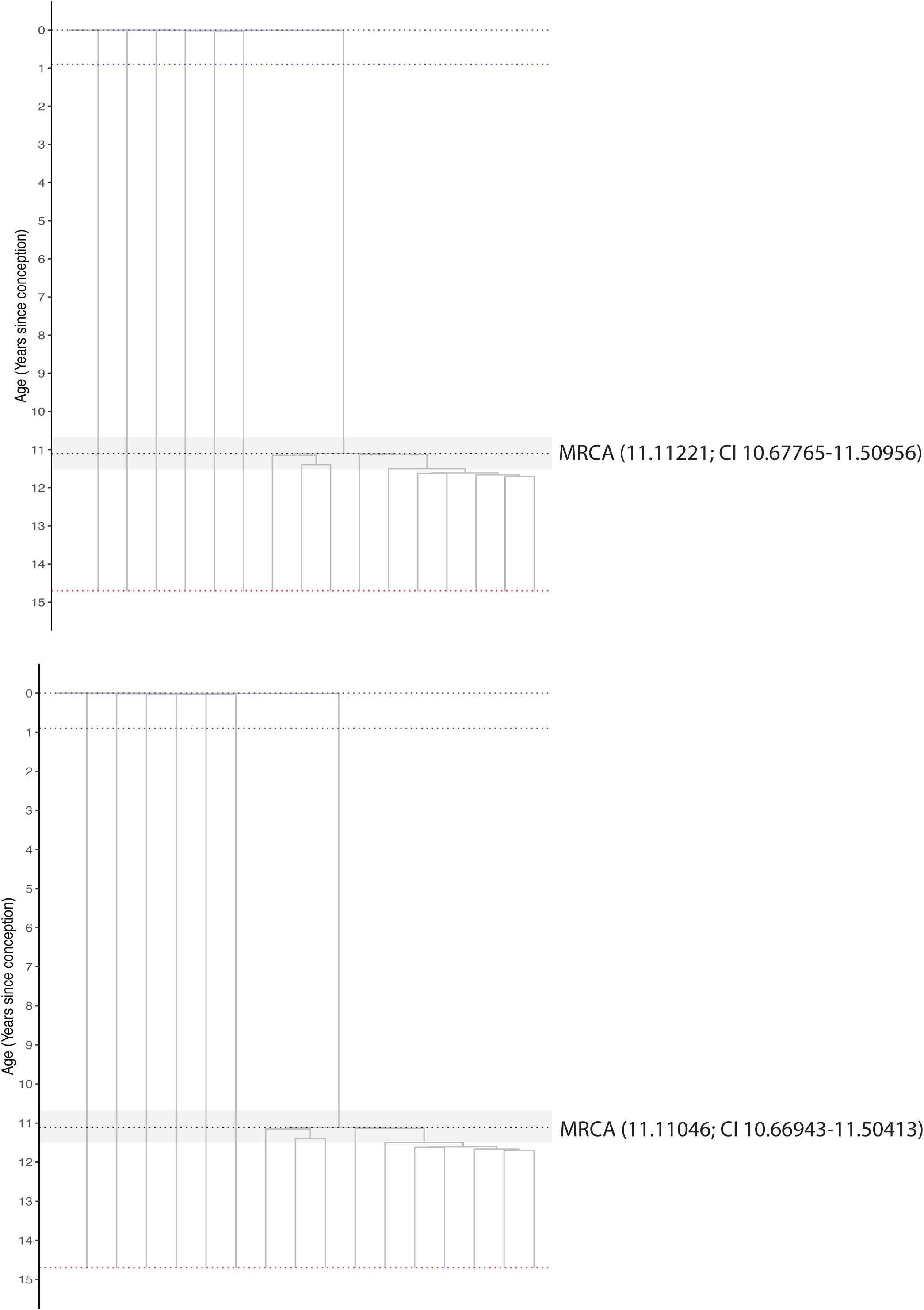
‘Time-based’ phylogenetic tree of patient P3G6, including all cells (top) and excluding one wild-type cell (P3G6GPDABC31) (bottom). Estimations of the emergence of the most recent common ancestor (MRCA) is shown on the right side of each tree, with 95% confidence intervals (CI).

